# VERSO: a comprehensive framework for the inference of robust phylogenies and the quantification of intra-host genomic diversity of viral samples

**DOI:** 10.1101/2020.04.22.044404

**Authors:** Daniele Ramazzotti, Fabrizio Angaroni, Davide Maspero, Carlo Gambacorti-Passerini, Marco Antoniotti, Alex Graudenzi, Rocco Piazza

## Abstract

We introduce VERSO, a two-step framework for the characterization of viral evolution from sequencing data of viral genomes, which improves over phylogenomic approaches for consensus sequences. VERSO exploits an efficient algorithmic strategy to return robust phylogenies from clonal variant profiles, also in conditions of sampling limitations. It then leverages variant frequency patterns to characterize the intra-host genomic diversity of samples, revealing undetected infection chains and pinpointing variants likely involved in homoplasies. On simulations, VERSO outperforms state-of-the-art tools for phylogenetic inference. Notably, the application to 6726 Amplicon and RNA-seq samples refines the estimation of SARS-CoV-2 evolution, while co-occurrence patterns of minor variants unveil undetected infection paths, which are validated with contact tracing data. Finally, the analysis of SARS-CoV-2 mutational landscape uncovers a temporal increase of overall genomic diversity, and highlights variants transiting from minor to clonal state and homoplastic variants, some of which falling on the spike gene. Available at: https://github.com/BIMIB-DISCo/VERSO.

## Introduction

The outbreak of coronavirus disease 2019 (COVID-19), which started in late 2019 in Wuhan (China)^1,2^ and was declared pandemic by the World Health Organization, is fueling the publication of an increasing number of studies aimed at exploiting the information provided by the viral genome of SARS-CoV-2 virus to identify its proximal origin, characterize the mode and timing of its evolution, as well as to define descriptive and predictive models of geographical spread and evaluate the related clinical impact^3–5^. As a matter of fact, the mutations that rapidly accumulate in the viral genome^6^ can be used to track the evolution of the virus and, accordingly, unravel the viral infection network^7,8^.

At the time of this writing, numerous independent laboratories around the world are isolating and sequencing SARS-CoV-2 samples and depositing them on public databases, e.g., GISAID^9^, whose data are accessible via the Nextstrain portal^10^. Such data can be employed to estimate models from genomic epidemiology and may serve, for instance, to estimate the proportion of undetected infected people by uncovering cryptic transmissions, as well as to predict likely trends in the number of infected, hospitalized, dead and recovered people^11–13^.

More in detail, most studies employ *phylogenomic approaches* that process *consensus sequences*, which represent the dominant virus lineage within each infected host. A growing plethora of methods for phylogenomic reconstruction is available to this end, all relying on different algorithmic frameworks, including distance-matrix, maximum parsimony, maximum likelihood or Bayesian inference, with various substitution models and distinct evolutionary assumptions (see, e.g.,^10,14–22^). However, while such methods have repeatedly proven effective in unraveling the main patterns of evolution of viral genomes with respect to many different diseases, including SARS-CoV-2^10, 23–25^, at least two issues can be raised.

First, most phylogenomics methods might produce unreliable results when dealing with *noisy* data, for instance due to sequencing issues, or with data collected with significant sampling limitations^14,26,27^, as witnessed for most countries during the epidemics^28,29^.

Second, most methods do not consider the key information on *intra-host minor variants* (also referred to as *minority variants* or *iSNVs*), which can be retrieved from whole-genome deep sequencing raw data and might be essential to improve the characterization of the infection dynamics and to pinpoint positively selected variants^30–32^. Due to the high replication, mutation and recombination rates of RNA viruses, subpopulations of mutant viruses, also known as viral *quasispecies*^30^, typically emerge and coexist within single hosts, and are supposed to underlie most of the adaptive potential to the immune system response and to anti-viral therapies^31,33,34^. In this regard, many recent studies highlighted the noteworthy amount of intra-host genomic diversity in SARS-CoV-2 samples^35–43^, similarly to what already observed in many distinct infectious diseases^8,32,44–48^.

Here, we introduce VERSO (Viral Evolution ReconStructiOn), a new comprehensive framework for the inference of high-resolution models of viral evolution from raw sequencing data of viral genomes (see Fig. 1). VERSO includes two consecutive algorithmic steps.

**Figure 1:**
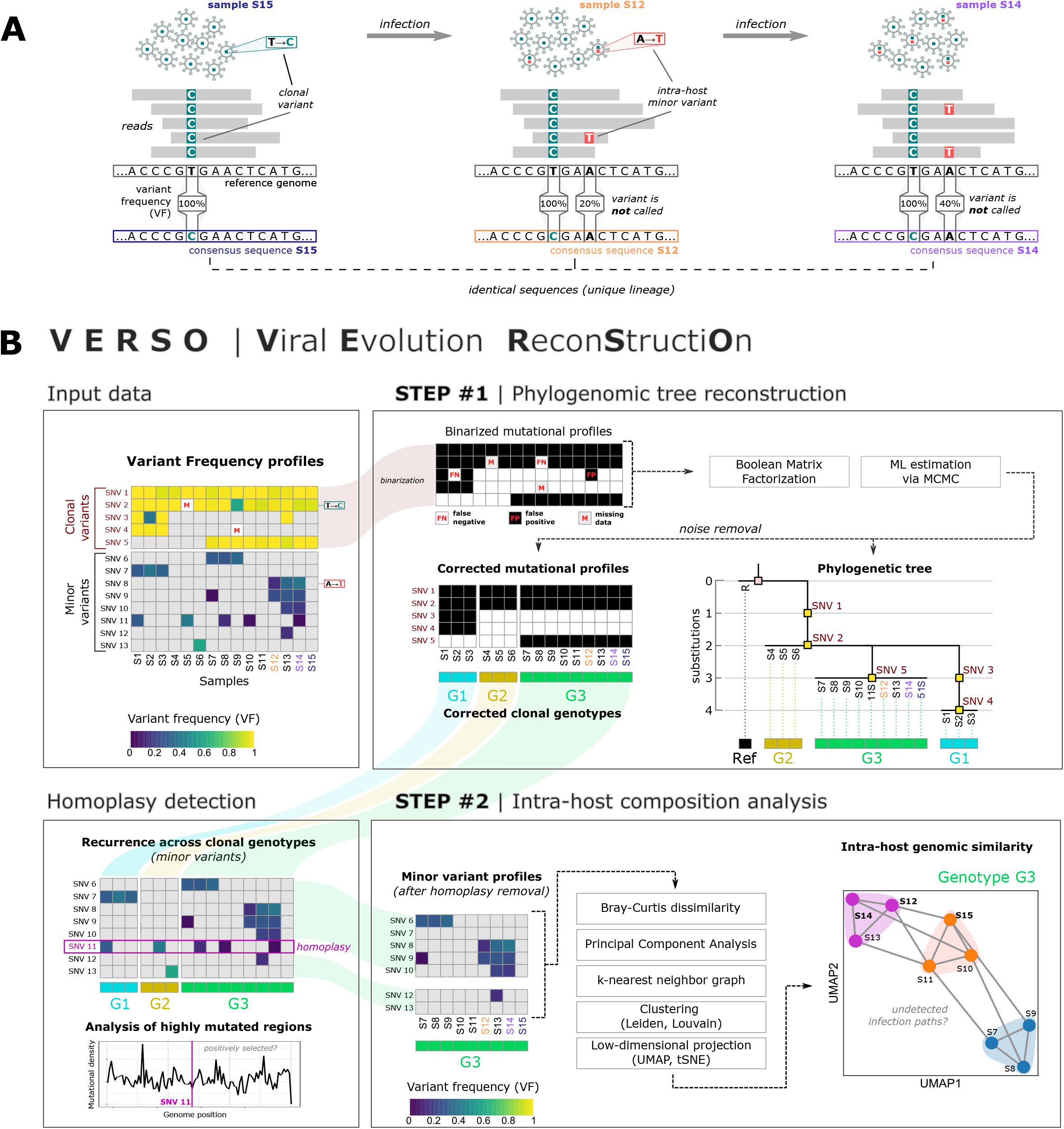
VERSO framework for viral evolution inference and intra-host genomic diversity quantification. **(A)** In this example, three hosts infected by the same viral lineage are sequenced. All hosts share the same clonal mutation (T>C, green), but two of them (#2 and #3) are characterized by a distinct minor mutation (A>T, red), which randomly emerged in host #2 and was transferred to host #3 during the infection. Standard sequencing experiments return an identical consensus sequence for all samples, by employing a threshold on variant frequency (VF) and by selecting mutations characterizing the dominant lineage. **(B)** VERSO takes as input the variant frequency profiles of samples, generated from raw sequencing data. In STEP #1, VERSO processes the binarized profiles of clonal variants and solves a Boolean Matrix Factorization problem by maximizing a likelihood function via Monte-Carlo Markov Chain, in order to correct false positives/negatives and missing data. As output, it returns both the corrected mutational profiles of samples and the phylogenetic tree, in which samples with identical corrected clonal genotypes are grouped in polytomies. Corrected clonal genotypes are then employed to identify homoplasies of minor variants, which are further investigated to pipoint positively selected mutations. The variant frequency profile of minor variants (excluding homoplasies) is processed by STEP #2 of VERSO, which computes a refined genomic distance among hosts (via Bray-Curtis dissimilarity on the kNN graph, after PCA) and performs clustering and dimensionality reduction, in order to project and visualize samples on a 2D space, representing the intra-host genomic diversity and the distance among hosts. This allows one to identify undetected transmission paths among samples with identical clonal genotype.

### STEP #1: Robust phylogenomic inference from clonal variant profiles

VERSO first employs a probabilistic noise-tolerant framework to process binarized clonal variant profiles (or, alternatively, consensus sequences), to return a robust phylogenetic model also in condition of sampling limitations and sequencing issues.

By adapting algorithmic strategies widely employed in cancer evolution analysis^49–52^, VERSO is able to correct false positive and false negative variants, can manage missing observations due to low coverage, and is designed to group samples with identical (corrected) clonal genotype in *polytomies*, avoiding ungrounded arbitrary orderings.

As a result, the accurate and robust phylogenomic models produces by VERSO may be used to improve the parameter estimation of epidemiological models, which typically rely on limited and inhomogeneous data^11,29^. Notice that this step can be executed independently from STEP #2, for instance in case raw sequencing data are not available.

#### Homoplasy detection (clonal variants)

The first step of VERSO allows one to identify clonal mutations that might be involved in *reticulation events*^53,54^ and, in particular, in *homoplasies*, possibly due to *positive selection* in a scenario of *convergent/parallel evolution*^55^, *founder effects*^31^ or *mutational hotspots*^56^. Such information might be useful to drive the design of opportune treatments and vaccines, for instance by blacklisting positively selected genomic regions.

### STEP #2: Characterization of intra-host genomic diversity

In the second step, VERSO exploits the information on *variant frequency* (VF) profiles obtained from raw-sequencing data (if available), to characterize and visualize the intra-host *genomic similarity* of hosts with identical (corrected) clonal genotype. In fact, even though the extent and modes of transmission of quasispecies from a host to another during infections are still elusive^31,57^, patterns of co-occurrence of minor variants detected in hosts with identical clonal genotype may provide an indication on the presence of undetected *infection paths*^8,58^.

For this reason, the second step of VERSO is designed to characterize and *visualize* the genomic similarity of samples by exploiting *dimensionality reduction* and *clustering* strategies typically employed in single-cell analyses^59^. Alternative approaches for the analysis of quasispecies, yet with different goals and algorithmic assumptions have been proposed, for instance in^60–63^ and recently reviewed in^64^.

As specified above, VERSO STEP #2 is executed on groups of samples with identical clonal genotype: the rationale is that the transmission of minor variants implicates the concurrent transfer of clonal variants, excluding the rare cases in which the VF of a clonal variant significantly decreases in a given host, for instance due to mutation losses (e.g., via recombination-associated deletions or via multiple mutations hitting an already mutated genome location^34^) or to complex horizontal evolution phenomena (e.g., super-infections^65,66^). Conversely, the transmission of clonal variants does not necessarily implicate the transfer of all minor variants, which are affected by complex recombination and transmission effects, such as bottlenecks^31,57^.

As a final result, VERSO allows one to visualize the genomic similarity of samples on a low-dimensional space (e.g., UMAP^67^ or tSNE^68^) representing the intra-host genomic diversity, and to characterize high-resolution infection chains, thus overcoming the limitations of methods relying on consensus sequences.

#### Homoplasy detection (minor variants)

Importantly, minor variants observed in hosts with distinct clonal genotypes (identified via VERSO STEP #1) may indicate homoplasies, due to mutational hotspots, phantom mutations or to positive selection^56^. VERSO pinpoints such variants for further investigations and allows one to exclude them from the computation of the VF-based genomic similarity prior to VERSO STEP #2, to reduce the possible confounding effects.

To summarize, VERSO: (*i*) returns accurate and robust phylogenies of viral samples, by removing noise from clonal variant profiles, (*ii*) detects reticulation events due to homoplasies of clonal variants, (*iii*) exploits minor variant profiles to characterize and visualize the intra-host genomic similarity of samples with identical (corrected) clonal genotype, thus pinpointing undetected infection paths, (*iv*) allows one to identify and characterize homoplastic minor variants, which might be due to positive selection or mutational hotspots.

To assess the accuracy and robustness of the results produced by VERSO, we performed an extensive array of simulations, and compared with two state-of-the-art methods for phylogenetic reconstruction, i.e., IQ-TREE^10^ and BEAST 2^22^. As a major result, VERSO outperforms competing methods in all settings and also in condition of high noise and sampling limitations.

Furthermore, we applied VERSO to two large-scale datasets, generated via Amplicon and RNA-seq Illumina sequencing protocols, including 3960 and 2766 samples, respectively. The robust phylogenomic models delivered via VERSO STEP #1 allow us to refine the current estimation on SARS-CoV-2 evolution and spread. Besides, thanks to the in-depth analysis of the mutational landscape of both clonal and minor variants, we could identify a number of variants undergoing transition to clonality, as well as several homoplasies, including variants likely undergoing positive selection processes.

Remarkably, the infection chains identified via VERSO STEP #2, by assessing the intra-host genomic similarity of samples with the same clonal genotype, were validated by employing contact tracing data from^69^. This important result, which could not be achieved by analyzing consensus sequences, proves the effectiveness of employing raw sequencing data to improve the characterization of the transmission dynamics, in particular during the early phase of the outbreak, in which a relatively low diversity of SARS-CoV-2 has been observed at the consensus level.

VERSO is released as free open source tool at this link:https://github.com/BIMIB-DISCo/VERSO.

## Results

### Comparative assessment on simulations

In order to assess the performance of VERSO and compare it with competing approaches, we executed extensive tests on simulated datasets, generated with the coalescent model simulator msprime^70^. Simulations allow one to compute a number of metrics with respect to the ground-truth, which in this case is the phylogeny of samples resulting from a backwards-in-time coalescent simulation^71^. Accordingly, this allows one to evaluate the accuracy and robustness of the results produced by competing methods in a variety of in-silico scenarios.

In detail, we selected 20 simulation scenarios with *n* = 1000 samples in which a number of clonal variants (with distinguishable profiles) between 14 and 31 was observed. We then inflated the datasets with false positives with rate *α* and false negatives with rate *β*, in order to mimic sequencing and coverage issues. Moreover, additional datasets were generated via random subsampling of the original datasets, to model possible sampling limitations and sampling biases. As a result, we investigated 4 simulations settings: (A) low noise, no subsampling, (B) high noise, no subsampling, (C) low noise, subsampling, and (D) high noise, subsampling (see Methods and the Supplementary Material for further details, the complete parameter settings of the simulations are provided in Supplementary Table S1).

VERSO STEP #1 was compared with two state-of-the-art phylogenetic methods from consensus sequences: IQ-TREE^10^, the algorithmic strategy included in the Nextstrain-Augur pipeline^72^, and BEAST 2^22^. Consensus sequences to be provided as input to such methods were generated from simulation data by employing the reference genome SARS-CoV-2-ANC (see below).

The performance of methods was assessed by comparing the reconstructed phylogeny with the simulated ground-truth, in terms of: (*i*) absolute error evolutionary distance, (*ii*) branch score difference^73^ and (*iii*) quadratic path difference^74^ (please refer to the Supplementary Material for a detailed description of all metrics).

In Figure 2 one can find the performance distribution of all methods with respect to all simulations settings. Notably, VERSO STEP #1 outperforms competing methods in all scenarios (Mann-Whitney U test *p <* 0.001 in all cases), with noteworthy percentage improvements, also in conditions of high noise and sampling limitations. This important result shows that the probabilistic framework that underlies VERSO STEP #1 can produce more robust and reliable results when processing noisy data, as typically observed in real-world scenarios.

**Figure 2:**
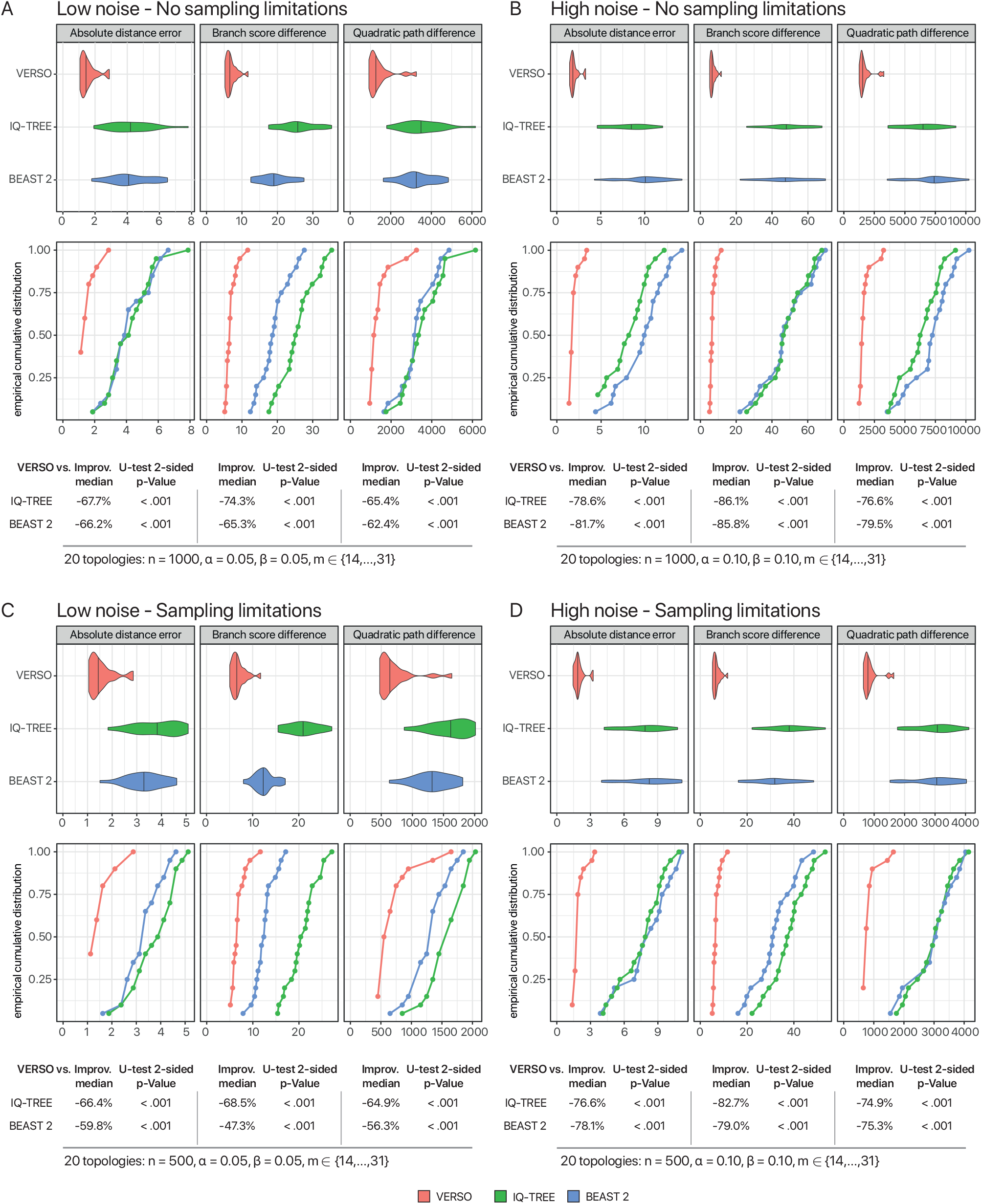
Comparative assessment on simulated data. Synthetic datasets were generated via the widely-used coalescent model simulator msprime^70^ – see the Supplementary Material and Table S1 of the Supplementary Material for the parameter settings. 20 distinct topologies with 1000 samples were generated including a number of distinguishable variants in the range (14, 31). For each topology, 4 synthetic datasets were generated, with different sample sizes (*n* = 1000, 500), and different combinations of false positives and false negatives ([*α* = 0.05, *β* = 0.05], [*α* = 0.10, *β* = 0.10]), for a total of 4 configurations and 80 independent datasets. VERSO STEP #1 was compared with IQ-TREE^10^ and BEAST 2^22^, on (*i*) absolute error evolutionary distance, (*ii*) branch score difference^73^ and (*iii*) quadratic path difference^74^ with respect to the ground-truth sample phylogeny provided by msprime (see the Supplementary Material for the description of the metrics). In the upper panels, distributions are shown as violin plots, whereas lower panels include the empirical cumulative distribution functions. The percentage improvement of VERSO with respect to competing methods is shown on all metrics (computed on median values), in addition to the p-value of the two-sided Mann-Whitney U test on distributions, for all settings.

### Reference genome

Different reference genomes have been employed in the analysis of SARS-CoV-2 origin and evolution. Two genome sequences from human samples, in particular, were used in early phylogenomic studies, namely sequence EPI_ISL_405839 (ref. #1 in the following) used, e.g., in^75^ and sequence EPI_ISL_402125 (ref. #2) used, e.g., in^3^. Excluding the polyA tails, the two sequences are identical for 29865 on 29870 genome positions (99.98%) and differ for only 5 SNPs at locations 8782, 9561, 15607, 28144 and 29095, for which ref. #1 has haplotype TTCCT and ref. #2 has haplotype CCTTC.

In order to define a likely common ancestor for both sequences, we analyzed the Bat-CoV-RaTG13 genome (sequence EPI_ISL_402131)^1^ and the Pangolin-CoV genome (sequence EPI_ISL_410721)^3,4^, which were identified as closely related genomes to SARS-CoV-2^76^. In particular, it was hypothesized that SARS-CoV-2 might be a recombinant of an ancestor of Pangolin-CoV and Bat-CoV-RaTG13^4, 77^, whereas more recent findings would suggest that SARS-CoV-2 lineage is the consequence of a direct or indirect zoonotic jump from bats^76^. Whatever the case, both Bat-CoV-RaTG13 and Pangolin-CoV display haplotype TCTCT at locations 8782, 9561, 15607, 28144 and 29095 and, therefore, one can hypothesize that such haplotype was present in the unknown common ancestor of ref. #1 and #2.

For this reason, we generated an artificial reference genome, named SARS-CoV-2-ANC, which is identical to both ref. #1 and #2 on 29865 (over 29870) genome locations, includes the polyA tail of ref. #2 (33 bases), and has haplotype TCTCT at locations 8782, 9561, 15607, 28144 and 29095 (see Supplementary Figure S2 for a depiction of the artificial genome generation). SARS-CoV-2-ANC is a likely common ancestor of both genomes and was used for variant calling in downstream analyses (SARS-CoV-2-ANC is released in FASTA format as Supplemetary File S1). Notice that VERSO pipeline is flexible and can employ any reference genome.

### Application of VERSO to 3960 samples from Amplicon sequencing data (Dataset #1)

We retrieved raw Illumina Amplicon sequencing data of 3960 SARS-CoV-2 samples of Dataset #1 and applied VERSO to the mutational profiles of 2906 samples selected after quality check (mutational profiles were generated by executing variant calling via standard practices; see Methods for further details). Notice that the analysis of this dataset was performed independently from that of Dataset #2 in order to exclude possible sequencing-related artifacts or idiosyncrasies.

### VERSO STEP #1 – Robust phylogenomic inference from clonal variant profiles

We first applied VERSO STEP #1 to the mutational profile of the 29 variants detected as clonal (VF > 90%) in at least 3% of the samples, in order to reconstruct a robust phylogenomic tree. The VERSO phylogenetic model is displayed in Fig. 3A and highlights the presence of 25 clonal genotypes, obtained by removing noise from data, and which define polytomies including different numbers of samples (see Methods for further details). The mapping between clonal genotype labels and the lineage dynamic nomenclature proposed in^78^ was obtained via pangolin 2.0^79^ and is provided in Supplementary File S3.

**Figure 3:**
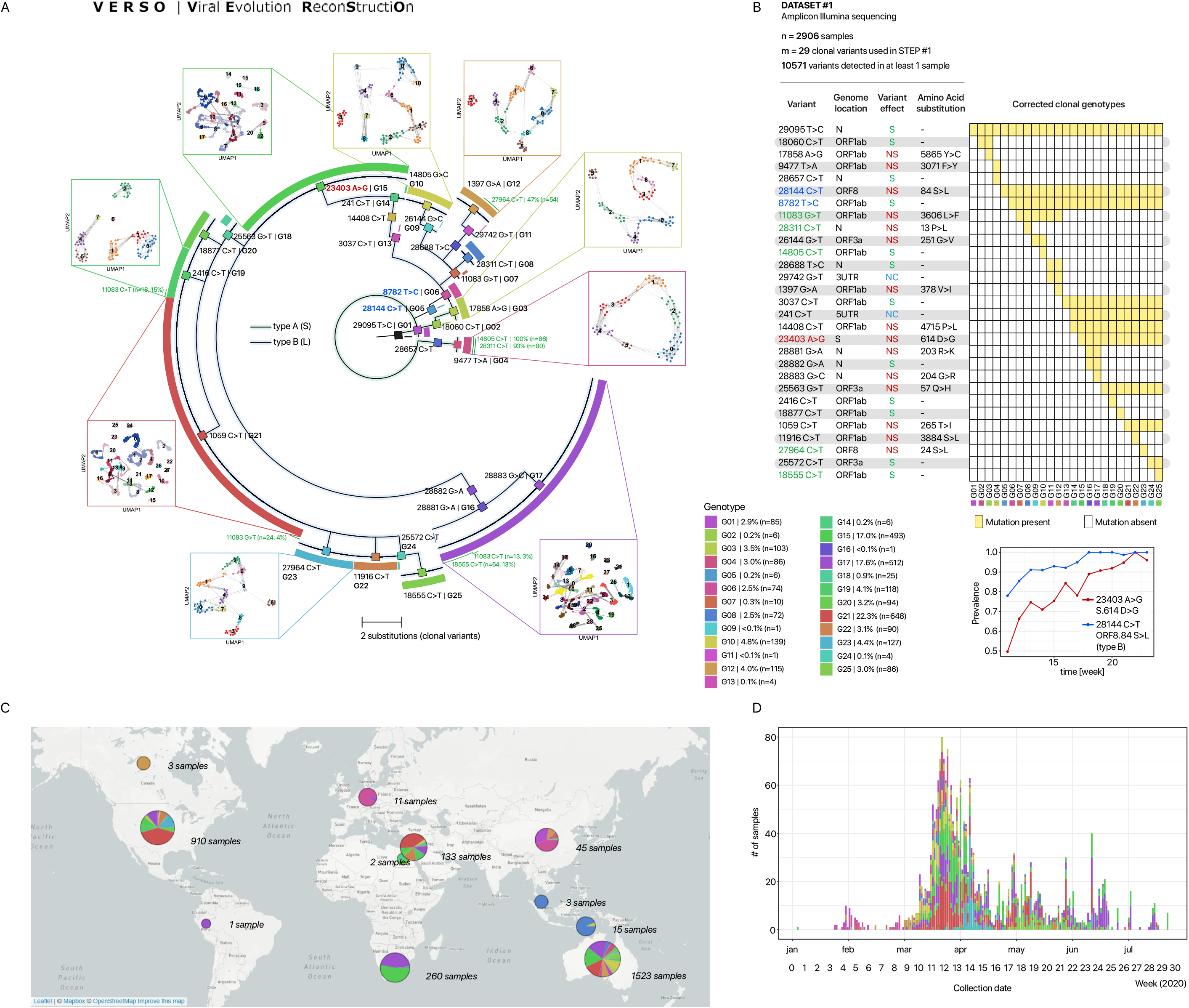
Viral evolution and intra-host genomic characterization of 2906 SARS-CoV-2 samples of via VERSO (Dataset #1). **(A)** The phylogetic model returned by VERSO STEP #1 from the mutational profile of 2906 samples selected after QC, on 29 clonal variants (VF > 90%) detected in at least 3% of the samples of Dataset #1 (reference genome: SARS-CoV-2-ANC). Colors mark the 25 distinct clonal genotypes identified by VERSO (the mapping with the lineage nomenclature proposed in^78^ and generated via pangolin 2.0^79^ is provided in Supplementary File S3). Samples with identical corrected clonal genotypes are grouped in polytomies and the black colored sample represents the SARS-CoV-2-ANC genome (visualization via FigTree^104^). The the green curves juxtaposed to certain polytomies report the number and fraction of samples in which the 5 homoplastic mutations are observed (only if the mutation is detected in at least 10 samples with the same corrected clonal genotype; see Supplementary File S2 for a summary on the samples exhibiting homoplastic clonal variants). The projection of the intra-host genomic diversity computed by VERSO STEP #2 from VF profiles is shown on the UMAP low-dimensional space for the clonal genotypes including *≥* 100 samples. Samples are clustered via Leiden algorithm on the k-nearest neighbor graph (*k* = 10), computed on the Bray–Curtis dissimilarity on VF profiles, after PCA. Solid lines represent the edges of the k-NNG. **(B)** The composition of the corrected clonal genotypes returned by VERSO STEP #1 is shown. Clonal SNVs are annotated with mapping on ORFs, synonymous (S), nonsynonymous (NS) and non-coding (NC) states, and related amino acid substitutions. Variants g.8782T>C (*ORF1ab*, synonymous) and g.28144C>T (*ORF8*, p.84S>L) are colored in blue, whereas variant g.23403 A>G (*S*, p.614 D>G) is colored in red. The prevalence variation in time of the relative haplotypes (i.e., the fraction of samples displaying such mutations) is also shown. The 5 homoplastic variants are colored in green. **(C)** The geo-temporal localization of the clonal genotypes via Microreact^87^ and **(D)** the prevalence variation in time are displayed.

More in detail, variant g.29095T>C (*N*, synonymous) is the earliest evolutionary event from reference genome SARS-CoV-2-ANC and is detected in 2454 samples of the dataset. The related clonal genotype G1, which is characterized by no further mutations, identifies a polytomy including 57 Australian, 15 Chinese, 12 American and 1 South-African samples.

Three clades originate from G1: a first clade includes clonal genotypes G2 (6 samples) and G3 (103), while a second clade includes clonal genotype G4 (86). Clonal genotypes G1–G4 are characterized by the absence of SNVs g.8782T>C (*ORF1ab*, synonymous) and g.28144C>T (*ORF8*, p.84S>L) and correspond to previously identified type A^24^ (also type S^80^), which was hypothesized to be an early SARS-CoV-2 type.

The third clade originating from clonal genotype G1 includes all remaining clonal genotypes (G5-G25) and is characterized by the presence of both SNVs g.8782T>C and g.28144C>T. This specific haplotype corresponds to type B^24^ (also type L^80^) and an increase of its prevalence has progressively recorded in the population, as one can see in Fig. 3, as opposed to type A (S), which was rarely observed in late samples. In this regard, we note that there are currently insufficient elements to support any epidemiological claim on virulence and pathogenicity of such SARS-CoV-2 types, even if recent evidences would suggest the existence of a low correlation^81^.

Variant g.23403A>G (*S*, p.614D>G) is of particular interest, as proven by the increasing number of related studies^82–85^. Such variant identifies a large clade including 11 clonal genotypes: G15 (493 samples), G16 (1), G17 (512), G18 (25), G19 (118), G20 (94), G21 (648), G22 (90),G23 (127), G24 (4), G25 (86), for a total of 2198 total samples, distributed especially in Australia (971), USA (841), South Africa (257) and Israel (125). Importantly, a constant increase of the prevalence of the haplotype corresponding to such variant is observed in time (see Fig. 3), which might hint at ongoing positive selection processes, e.g., due to increased viral transmission. However, this hypothesis is highly debated^86^ and, in order to investigate the possible functional effect of such variant and the related clinical implications, *in vivo* and *in vitro* studies are needed^85^.

By looking more in detail at the geo-temporal localization of samples depicted via Microreact^87^ (Figure 3B), one can see that the different clonal genotypes are distributed across the world in distinct complex patterns, suggesting that most countries might have suffered from multiple introductions, especially in the early phases of the epidemics. In particular, samples are distributed in 11 countries, with Australia (1523 samples), USA (910), South Africa (260), Israel (133) and China (45) representing around 99% of the dataset.

The country displaying the largest number of samples is Australia, with 1523 samples, distributed in 22 different clonal genotypes. The presence of a number of early clonal genotypes (i.e. G1, G2, G3, G4 and G6) supports the hypothesis of multiple introductions of SARS-CoV-2 in Australia. Interestingly, we note that from the 16^th^ week on the composition of the Australian sample group tends to be polarized towards clonal genotypes G17 (108*/*311 ≈ 35%) and G25 (82*/*311 ≈ 26%).

910 samples from USA are included in the dataset, distributed in 17 different clonal genotypes, with G21 being the most abundant in the population (376*/*910 ≈ 41%). Also in this case, samples collected in the initial weeks belong to the ancestral clades, supporting the hypothesis of multiple introductions. Notably, after the 17^th^ week all American samples display the haplotype g.23403A>G (*S*,p.614D>G) and we notice an overall decrease in genomic diversity, since the observed clonal genotypes pass from 16 (week interval 9 −16, 2020) to 8 (week interval 17 −29, 2020). Notice that only 49.1% of the American samples have a collection date.

260 samples from South Africa are included in the dataset, which are partitioned in 6 different clonal genotypes, 4 of which (G1, G8, G14 and G16) include a single sample, whereas 98.46% of the samples exhibit the haplotype g.23403A>G (*S*,p.614D>G) and, specifically, are included in clonal genotypes G15 and G17. Finally, all Chinese samples were collected in the early phase (January–February, 2020) and are characterized by 6 different clonal genotypes (i.e., G1, G6, G7, G9, G11 and G12).

#### Homoplasy detection (clonal variants

5 clonal variants included in our model show apparent violations of the accumulation hypothesis, namely: g.11083G>T (*ORF1ab*, p.3606 L>F), g.14805C>T (*ORF1ab*, synonymous), g.18555C>T (*ORF1ab*, synonymous), g.27964C>T (*ORF8*, p.24S>L), and g.28311C>T (*N*,p.13P>L), suggesting that they might be involved in homomplasies. In Figure 3 the samples in which the 5 homoplastic variants are detected are highlighted (if the mutation is detected in ≥ 10 samples with the same corrected clonal genotype), whereas in Supplementary Figure S3 one can find the expanded clonal variant tree, in which the reticulation related to such variants is explicitly depicted.

Some of such variants have been exhaustively studied (e.g., g.11083G>T in^88^), specifically to verify possible scenarios of convergent evolution, which may unveil the fingerprint of adaptation of SARS-CoV-2 to human hosts. To this end, particular attention should be devoted to the three non synonymous substitutions, i.e., g.11083G>T (present in 460 samples, ≈16% of the dataset), g.27964C>T (182 samples, ≈ 6%) and g.28311C>T (153 samples, ≈5%). As a first result, we note the prevalence dynamics of the haplotypes defined by such variants does not show any apparent growth trend in the population (see Supplementary Figure S5).

To further investigate if such variants fall in a region prone to mutations of the SARS-CoV-2 genome, we evaluated the mutational density employing a sliding window approach similarly to^89^ (see Supplementary Material for additional details). As shown in Supplementary Figure S4, the mutational density, computed by considering synonymous minor variants, exhibits a median value of = 0.083 [syn.mutations][nucleotides]^*−*1^. Interestingly, the three nonsynonymous SNVs (g.11083G>T, g.27964C>T and g.28311C>T) are located within windows with a higher mutational density than the median value: 0.085, 0.124 and 0.1 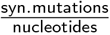, respectively (see Table S5 of the Supplementary Material), and this would suggest that they might have originally emerged due to the presence of natural mutational hotspots or phantom mutations.

However, this analysis is not conclusive and further investigations are needed to characterize the functional effect of such mutations and the possible impact in the evolutionary and diffusion process of SARS-CoV-2.

##### Stability analysis

The choice of an appropriate VF threshold to identify clonal variants and, accordingly, to generate consensus sequences from raw sequencing data might affect the stability of the results of any downstream phylogenomic analysis. On the one hand, loose thresholds might increase the risk of including non-clonal variants in consensus sequences. On the other hand, too strict thresholds might increase the rate of false negatives, especially with noisy sequencing data.

For this reason, we assessed the robustness of the results produced by VERSO STEP #1 on Dataset #1 when different thresholds in the set *δ* ∈{0.5, 0.6, 0.7, 0.8} are employed to identify clonal variants, with those obtained with default threshold (*δ* = 0.9), in terms of tree accuracy (see the Supplementary Material for further details). As one can see in Supplementary Figure S7, the tree accuracy varies between 0.97 and 0.98 in all settings, proving the the results produced by VERSO STEP #1 are robust with regard to the choice of the VF threshold for clonal variant identification.

#### VERSO STEP #2 – Characterization of intra-host genomic diversity

We then applied VERSO STEP #2 to the complete VF profiles of the samples with the same clonal genotype and projected their intra-host genomic diversity on the UMAP low-dimensional space. This was done excluding: (*i*) the clonal variants employed in the phylogenetic inference via VERSO STEP #1, (*ii*) all minor variants (VF ≤90%) observed in more than one clonal genotype (i.e., homoplasies) and which are likely emerged independently within the hosts, due to mutational hotspots, phantom mutations or positive selection (see Methods and the next subsections). Even though, as expected, the VF profiles of minor variants are noisy, a complex intra-host genomic architecture is observed in several individuals. Moreover, patterns of co-occurrence of minor variants across samples support the hypothesis of transmission from a host to another.

In Fig. 3 we display the UMAP plots for the clonal genotypes including more than 100 samples, plus clonal genotype G4 (*n* = 86 samples), which was used for contact tracing analyses. Such maps describe likely transmission paths among hosts characterized by the same (corrected) clonal genotype and, in most cases, suggests the existence of several distinct infection clusters with different size and density. This result was achieved by exploiting the different properties of clonal and minor variants via the two-step procedure of VERSO.

##### Contact tracing

To corroborate our findings, we employed the contact tracing data from^69^, in which 65 samples from Dataset #1 are characterized with respect to household, work location or other direct contacts. 4 distinct contact groups, including 36, 15, 12 and 2 samples, respectively, are associated directly or indirectly to three different New South Wales institutions (i.e., Institutions #1, #2 and #3) and to the same household environment (Household #1).

As a first result, all samples belonging to a specific contact group are characterized by the same clonal genotype, determined via VERSO STEP #1, a result that confirms recent findings^42,69^. More importantly, the analysis of the intra-host genomic diversity via VERSO STEP #2 allows one to highly refine this analysis.

In Fig. 4 one can find the UMAP plot of clonal genotypes G4, G12 and G21, which include 36 (over 86), 14 (over 115) and 14 (over 648) samples with contact information. Strikingly, the distribution of the pairwise intra-host genomic distance among samples from the same institution/household (computed on the k-nearest neighbour graph via Bray-Curtis dissimilarity, after PCA; see Methods) is significantly lower with respect to the distance of all samples with the same clonal genotype (p-value of the Mann-Whitney U test *<* 0.001 in all cases). Furthermore, all samples belonging to the same contact group are connected in the kNN graph, while a noteworthy proportion of samples without contact information in genotypes G12 and G21 are placed in disconnected graphs (24.9% and 76.4%, respectively).

**Figure 4:**
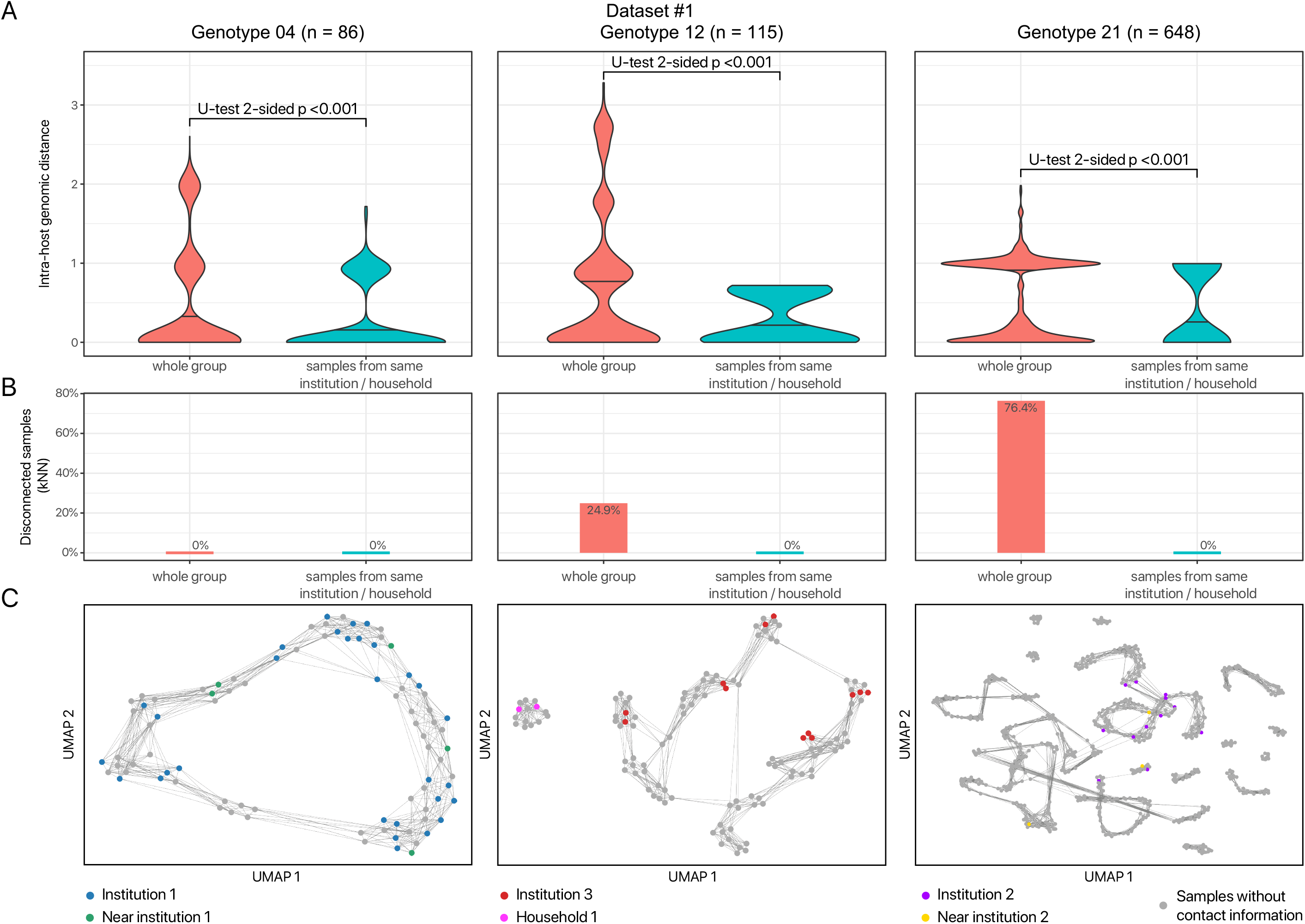
Infection dynamics revealed via characterization of intra-host genomic similarity – Dataset #1. **(A)** The distribution of the pairwise intra-host genomic distance (computed via Bray-Curtis dissimilarity on the kNN graph, wih *k* = 10, after PCA; see Methods) for the samples belonging to the same household or institution (including samples marked as near), versus the pairwise distance of all samples belonging to clonal genotypes G4, G12 and G21. P-values of the Mann-Whitney U test two-sided are also shown. **(B)** The proportion of samples that are disconnected in the kNN graph, with respect to the samples belonging to the same household or institution (including samples marked as near) and with respect to all samples. **(C)** The UMAP projection of the intra-host genomic diversity of the samples belonging to clonal genotypes G4, G12 and G21, returned by VERSO STEP #2.

This major result suggests that patterns of co-occurrence of minor variants can indeed provide useful indication on contact tracing dynamics, which would be masked when employing consensus sequencing data. Accordingly, the algorithmic strategy employed by VERSO STEP #2 and, especially, the identification of the kNN graph on intra-host genomic similarity, provides an effective tool to dissect the complexity of viral evolution and transmission, which might in turn improve the reliability of currently available contact tracing tools.

##### Homoplasy detection (minor variants)

Several minor variants are found in samples with distinct clonal genotypes and might indicate the presence of homoplasies. In this respect, the heatmap in Fig. 5F returns the distribution of minor SNVs with respect to: (*i*) the number of distinct clonal genotypes in which they are detected and (*ii*) the mutational density of the region in which they are located (see the Supplementary Material for details on the mutational density analysis).

**Figure 5:**
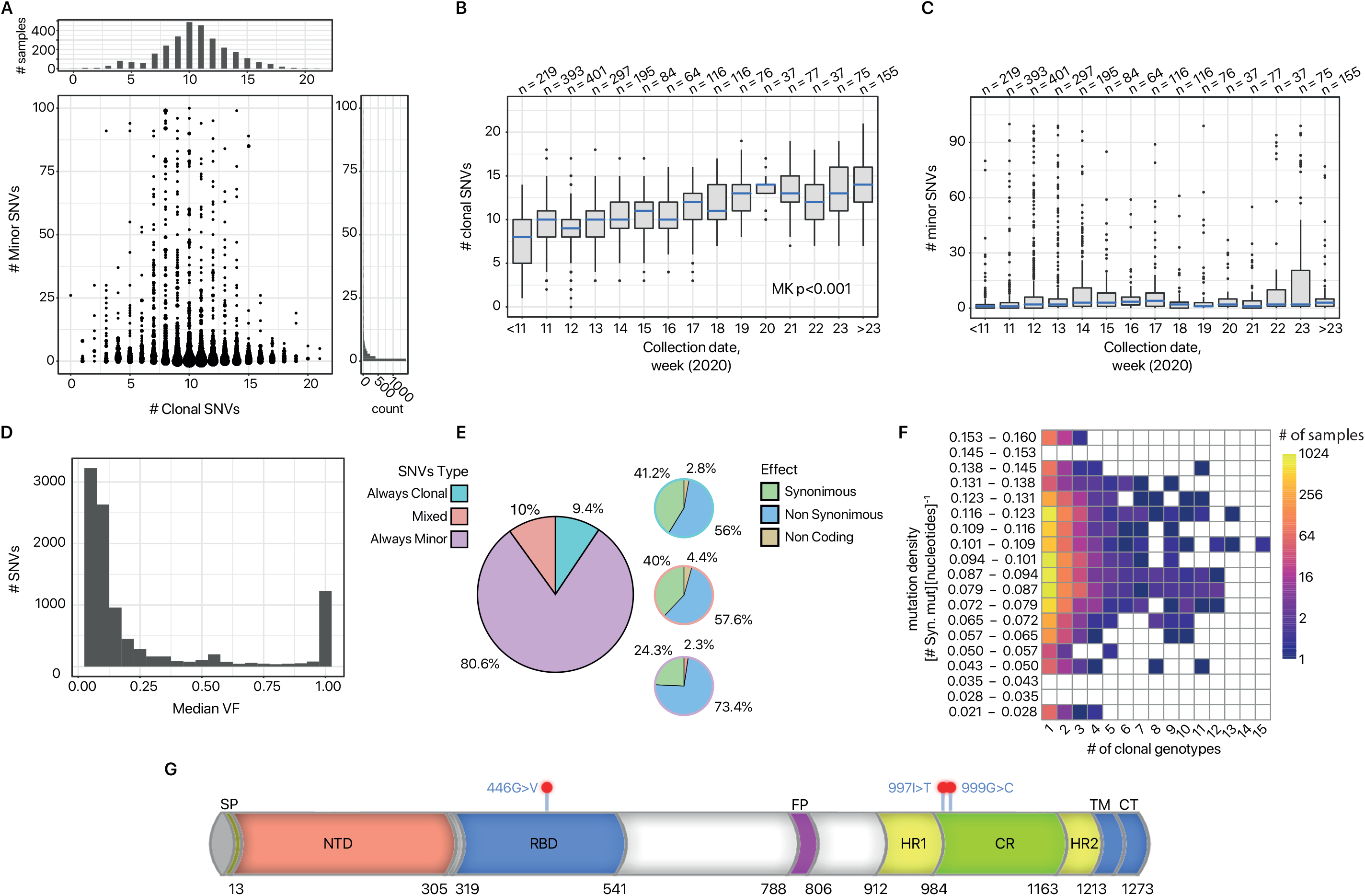
Mutational landscape of 2906 SARS-CoV-2 samples (Dataset #1). **(A)** Scatterplot displaying, for each sample, the number of clonal (VF > 90%) and minor variants (VF *≤* 90%, node size proportional to the number of samples). **(B-C)** Boxplots returning the distribution of the number of clonal and minor variants, obtained by grouping samples according to collection date (weeks, 2020). The p-value of the Mann-Kendall (MK) trend test on clonal variants is highly significant. **(D)** Distribution of the median VF for all SNVs detected in the viral populations. **(E)** Pie-charts returning: (left) the proportion of SNVs detected as always clonal, always minor or mixed; (right), for each category, the proportion of synonymous, nonsynonymous and non coding variants (check the pie-chart border color for a visual clue). **(F)** Heatmap returning the distribution of always minor SNVs with respect: (x axis) the number of clonal genotype of the phylogenonic model in Fig. 3 in which each variant is observed, (y axis) the mutational density of the genome region in which it is located (see the Supplementary Material) **(G)** Mapping of the candidate homoplastic minor variants located on the spike gene of the SARS-CoV-2 virus.

The intuition is that the variants detected in single clonal genotypes (left region of the heatmap) are likely spontaneously emerged private mutations, or the result of infection events between hosts with same clonal genotype (see above). Conversely, SNVs found in multiple clonal genotypes (right region of the heatmap) may have emerged due to positive selection in a parallel/convergent evolution scenario, or to mutational hotspots or phantom mutations. To this end, the mutational density analysis provides useful information to pinpoint mutation-prone regions of the genome.

Interestingly, a significant number of minor variants are observed in multiple clonal genotypes and fall in scarcely mutated regions of the genome (see Supplementary Figure S4). This would suggest that some of these variants might have been positively selected, due to some possible functional advantage or to transmission-related founder effects. In this respect, we further focused our investigation on a list of 80 candidate minor variants, which: (*i*) are detected in more than one clonal genotype, (*ii*) are present in at least 10 samples, (*iii*) are nonsynonymous and (*iv*) fall in a region of the genome with mutational density lower than the median value (see Table S6 of the Supplementary Material for details on such variants). In the following, we focus on a subset of such variants falling on the spike gene of the SARS-CoV-2 genome.

##### Considerations on homoplasies falling on the spike gene

The spike protein of SARS-CoV-2 plays a critical role in the recognition of the ACE2 receptor and in the ensuing cell membrane fusion process^90^. We prioritized 3 candidate homoplastic minor variants occurring on the SARS-CoV-2 spike gene (S) (see Table S6 of the Supplementary Material). Interestingly, 2 out of 3, namely: g.24552T>C (p.997I>T) and g.24557G>T (p.999G>C), detected in 57 samples in total (10 and 47 samples, respectively), clustered in the so-called Connector Region (CR), bridging between the two heptad repeat regions (HR1 and HR2) of the S2 subunit of the spike protein.

When the Receptor Binding Domain (RBD) binds to ACE2 receptor on the target cell, it causes a conformational change responsible for the insertion of the Fusion Peptide (FP) into the target cell membrane. This, in turn, triggers further conformational changes eventually promoting a direct interaction between HR1 trimer and HR2, which occurs upon bending of the flexible CR, in order to form a six-helical HR1-HR2 complex known as the Fusion Core Region (FCR) in close proximity to the target cell plasma membrane, ultimately leading to viral fusion and cell entry^91^.

Peptides derived from the HR2 heptad region of enveloped viruses and able to efficiently bind to the viral HR1 region inhibit the formation of the FCR and completely suppress viral infection^92^. Therefore, the formation of the FCR is considered to be vital to mediate virus entry in the target cells, promoting viral infectivity. Of note, the CR is highly conserved across the gammacoronavirus genus, supporting the notion that this region may play a very important but yet unclear functional role (Fig 5G). Although structural and in vitro models will be required in order to extensively characterize the functional effect of these variants, the evidence that two our of three minor variants detected in the spike protein falls in a small domain comprising less than 14% of the entire spike protein length is intriguing, as it suggests a potential functional role for these mutations. It will be important to track the prevalence of these mutations, as well as of all other candidate convergent variants falling on different region of the SARS-CoV-2, to highlight possible transitions to clonality (see below). We also remark that, being a data science computational approach, VERSO can struggle in dissecting complex mutational cases, since all the experimental hypotheses that can be generated are clearly data-dependent. For this reason, and given the heterogeneity and limitations of currently available SARS-CoV-2 datasets, any hypothesis delivered by VERSO requires additional independent investigations and ad hoc experimental validations.

#### Mutational landscape

We analyzed in-depth the mutational landscape of the samples of Dataset #1. First, the comparison of the number of clonal (VF > 90%) and minor variants detected in each host (Fig. 5A) reveals a bimodal distribution of clonal variants (with first mode at 4 and second mode at 10), whereas minor variants display a more dispersed long-tailed distribution with median equal to 2 and average ≈23. From the plot, it is also clear that individuals characterized by the same clonal genotype may display a significantly different number of minor variants, with distinct distributions observed across clonal genotypes.

The comparison of the distribution of the number of variants obtained by grouping the samples with respect to collection week (Fig. 5B-C) allows us to highlight a highly statistically significant increasing trend for clonal variants (Mann-Kendall trend test on median number of clonal variants *p <* 0.001). This result would strongly support both the hypotheses of accumulation of clonal variants in the population and that of a concurrent increase of overall genomic diversity of SARS-CoV-2^36, 93^, whereas the relevance of this phenomenon on minor variants is unclear.

We then focused on the properties of the SNVs detected in the population. Surprisingly, the distribution of the median VF for each detected variants (Fig. 5D) reveals a bimodal distribution, with the large majority of variants showing either a very low or a very high VF, with only a small proportion of variants showing a median VF within the range 10 −90%. This behavior is typical of systems where the prevalence of some subpopulations is driven by positive Darwinian selection while others are purified^94^.

In order to analyze the two components of this distribution, we further categorized the variants as *always clonal*, (i.e., SNVs detected with VF > 90% in all samples), *always minor* (i.e., SNVs detected with VF 5% and ≤ 90% in all samples) and *mixed* (i.e., SNVs detected as clonal in at least one sample and as minor in at least another sample). As one can see in Fig. 5E, 9.4%, 80.6% of and 10% all SNVs are respectively detected as always clonal, always minor and mixed in our dataset. Moreover, 56%, 73.4% and 57.6% of always clonal, always minor and mixed variants, respectively, are nonsynonymous, whereas the large majority of remaining variants are synonymous.

These results would suggest that, in most cases, randomly emerging SARS-CoV-2 minor variants tend to remain at a low frequency in the population, whereas, in some circumstances, certain variants can undergo frequency increases and even become clonal, due to undetected mixed transmission events or to selection shifts, as it was observed in^8^ for the cases of H3N2 and H1N1/2009 influenza. Interestingly, 15 variants identified as possibly convergent (see above) fall in this category and deserves further investigations (see Table S6 of the Supplementary Material for additional details).

##### Transmission bottleneck analysis

The estimation of transmission bottlenecks might be of specific interest during the current pandemics. Despite most available methods require data collected on donor-host couples (see, e.g.,^95,96^), here we employed a strategy akin to^97,98^ and that is roughly based on the analysis of the variation of the VF variance of a number of candidate neutral mutations. The intuition is that variance shrinking indicates significant transmission bottlenecks which, accordingly, would result in lower viral diversity transferred from a host to another and, possibly, in purification of certain variants in the population. As the analysis ideally requires the comparison of groups in which infection events have occurred, here we considered groups of samples with distinct clonal genotypes, separately. We then selected a number of variants as neutral markers. The rationale is that transmission phenomena such as bottlenecks are expected to significantly affect the VF variance of neutral markers (please see Supplementary Material for further details).

More in detail, we first split the samples of each clonal genotype, for which a collection date is available, in non-overlapping groups corresponding to two consecutive time windows, i.e., before and after the 14^th^ week, 2020. Accordingly, 3 SNVs were selected as candidate neutral or quasi-neutral markers, namely variants g.634 T>C, g.14523 A>G and g.15168 G>A. In Supplementary Figure S6, one can find the distribution of the variant frequency of the selected markers with respect to the time windows, which highlights moderate variations of the variance for all markers (see also Table S7 of the Supplementary Material). All in all, this result would suggest the presence of mild bottleneck effects, consistently with recent studies involving donor-host data^43^.

#### Application of VERSO to 2766 samples from RNA-sequencing data (Dataset #2)

We retrieved the raw Illumina RNA-sequencing data of 2766 samples included in Dataset #2 and applied VERSO to the mutational profiles of 1438 samples selected after quality check. 23 clonal variants were employed in the analysis, according to the filters described in the Methods Section.

The resulting phylogenetic model is consistent with the one obtained for Dataset #1, despite minor differences (Supplementary Figure S8A). Specifically, 18 distinct clonal genotypes are identified by VERSO STEP #1, 11 of which are identical to those found in the analysis of Dataset #1 (in such cases the same genotype label was maintained; see Supplementary File S3 for the mapping with the lineage nomenclature proposed in^78^). 5 further clonal genotypes are evolutionary consistent and represent independent branches detected due to the non-overlapping composition of the dataset, and are labeled with progressive letters from the closest genotype (i.e., G13a, G21a, G22a, G22b, G23a), while the 2 samples of genotype G13a* might be safely assigned to genotype G13a, since the absence of mutation g.3037C>T is likely due to low coverage.

By excluding the remaining clonal genotype GH, which presents inconsistencies due to the presence of the candidate homoplastic variant g.11083G>T (*ORF1ab*, p.3606L>F, see above), all clonal genotypes display the same ordering in both Datasets (also see the expanded clonal variant tree in Supplementary Figure S9). This proves the robustness of the results delivered by VERSO STEP #1 even when dealing with data generated from distinct sequencing platforms.

By looking at the geo-temporal localization of samples obtained via Microreact^87^ (Supplementary Figure S8B), one can see that that Dataset #2 includes samples with a significantly different geographical distribution with respect to Dataset #1. This dataset contains sample from 10 countries, with the large majority collected in USA (96.8%). More in detail, the samples of such country are mostly characterized by clonal genotype G21. We further notice that, also for Dataset #2, mutation g.23403A>G (*S*,p.614D>G) becomes prevalent in the population at late collection dates. Moreover, only samples belonging to previously defined type B are detected in this dataset.

The analysis of the intra-host genomic diversity was also performed for Dataset #2 via VERSO STEP #2, which would suggest the existence of undetected infection events and of several infection clusters with distinct properties, even though no contact tracing are available in this case. Overall, this proves the general applicability of the VERSO framework, which can produce meaningful results when applied to data produced with any sequencing platforms. However, in order to minimize the possible impact of data- and platform-specific biases, we suggest to perform the VERSO analysis on datasets generated from different protocols separately.

#### Scalability

We finally assessed the computational time required by VERSO in a variety of simulated scenarios. The results are shown in the Supplementary Material (Supplementary Figure S10) and demonstrate the scalability of VERSO also when processing large-scale datasets.

## Discussion

We introduced VERSO, a comprehensive framework for the high-resolution characterization of viral evolution from sequencing data, which improves over currently available methods for the analysis of consensus sequences. VERSO exploits the distinct properties of clonal and minor variants to dissect the complex interplay of genomic evolution *within* hosts and transmission *among* hosts.

On the one hand, the probabilistic framework underlying VERSO STEP #1 delivers highly accurate and robust phylogenetic models from clonal variants, also in condition of noisy observations and sampling limitations, as proven by extensive simulations and by the application to two-large scale SARS-CoV-2 datasets generated from distinct sequencing platforms. On the other hand, the characterization of intra-host genomic diversity provided by VERSO STEP #2 allows one to identify undetected infection paths, which were in our case validated with contact tracing data, as well as to intercept variants involved in homoplasies.

This may represents a major advancement in the analysis of viral evolution and spread and should be quickly implemented in combination to data-driven epidemiological models, to deliver a high-precision platform for pathogen detection and surveillance^12,99^. This might be particularly relevant for countries which suffered outbreaks of exceptional proportions and for which the limitations and inhomogeneity of diagnostic tests have proved insufficient to define reliable descriptive/predictive models of disease diffusion. For instance, it was hypothesized that the rapid diffusion of COVID-19 might be likely due to the extremely high number of untested asymptomatic hosts^100^.

More accurate and robust phylogenetic models may allow one to improve the assessment of molecular clocks and, accordingly, the estimation of the parameters of epidemiological models such as SIR and SIS^11,101^, as well as to unravel the cryptic transmission paths^8,12,13,102^. Furthermore, the finer grain of the analysis on intra-host genomic similarity from sequencing data might be employed to enhance the active surveillance, for instance by facilitating the identification of infection clusters and super-spreaders^103^. Finally, the characterization of variants possibly involved in positive selection processes might be used to drive the experimental research on treatments and vaccines.

## Experimental Procedures

### Resource Availability

#### Lead Contact

Alex Graudenzi, Institute of Molecular Bioimaging and Physiology, Consiglio Nazionale delle Ricerche (IBFM-CNR), via F.lli Cervi, 93, 20090 Segrate, Milan, Italy.

#### Materials Availability

This study did not generate new unique reagents.

#### Data and Code Availability

VERSO is freely available at this link: https://github.com/BIMIB-DISCo/VERSO. VERSO STEP #1 is provided as an open source standalone R tool, whereas STEP #2 is provided as a Python script. The source code to replicate all the analyses presented in the manuscript, both on simulated and real-world datasets, is available at this link: https://github.com/BIMIB-DISCo/VERSO-UTILITIES.

SCANPY^59^ is available at this link: https://scanpy.readthedocs.io/en/stable/. The Web-based tool for the geo-temporal visualization of samples, Microreact^87^, is available at this link: https://microreact.org/showcase. The tool employed to plot the phylogenomic model returned by VERSO STEP #1 (in Newick file format) is FigTree^104^ and is available at this link: http://tree.bio.ed.ac.uk/software/figtree/. The tool used for the mapping between clonal genotype labels and the dynamic nomenclature proposed in^78^ is pangolin 2.0^79^ and is available at this link: https://github.com/cov-lineages/pangolin.

### VERSO STEP #1: robust phylogenomic inference from clonal variant profiles

VERSO is a novel framework for the reconstruction of viral evolution models from raw sequencing data of viral genomes. It includes a two-step procedure, which we describe in the following.

The first step of VERSO employs a probabilistic maximum-likelihood framework for the reconstruction of robust phylogenetic trees from binarized mutational profiles of clonal variants (or, alternatively, from consensus sequences). This step relies on an evolved version of the algorithmic framework introduced in^105^ for the inference of cancer evolution models from single-cell sequencing data, and can be executed independently from STEP #2, in case raw sequencing data are not available.

#### Inputs

The method takes as input a *n* (samples) ×*m* (variants) binary mutational profile matrix, as defined on the basis of clonal SNVs only. In this case, an entry in a given sample is equal to 1 (present) if the VF is larger than a certain threshold (in our analyses, equal to 90%), it is equal to 0 if lower than a distinct threshold (in our analyses, equal to 5%), and is considered as missing (NA) in the other cases, thus modeling possible uncertainty in sequencing data or low coverage.

Notice that consensus sequences can be processed by VERSO STEP #1 by generating a consistent binarized mutational profile matrix. We also note that given the intrinsic challenges associated with a reliable identification of low VF indels, the analysis focuses only on single nucleotide variants. Further details on the variant calling pipeline employed in this study are provided in the next subsections.

#### The algorithmic framework

VERSO STEP #1 is a probabilistic framework which solves a Boolean matrix factorization problem with perfect phylogeny constraints and relying on the Infinite Sites Assumption (ISA)^106,107^. The ISA subsumes a consistent process of accumulation of clonal variants characterizing the evolutionary history of the virus and does not allow for losses of mutations or convergent variants (i.e., mutations observed in distinct clades).

In this regard, we recall that that the variant accumulation hypothesis holds only when considering clonal mutations. In fact, clonal mutations (e.g., A, B, C, D and E) are present – by definition – in the large majority of the quasispecies of a given sample, depending on the chosen VF threshold (in our case equal to 90%, see above). Since such variants are rarely lost, they are most likely transmitted from a host to another during infections. In addition, the origination of new clonal mutations in single samples leads to the definition of new clonal genotypes, following a standard branching process (e.g., A, AB, ABC, ABD, ABDE). As a result, clonal mutations typically accumulate during the evolutionary history of a virus, excluding complex scenarios involving reticulation events^53^, whereas clonal genotypes can clearly get extinct. Conversely, variants with lower frequency do not necessarily accumulate, due to the high recombination rates, as well as to bottlenecks, founder effects and stochasticity^31^, and this is the reason why they were considered separately in the analysis, via VERSO STEP #2 (see below).

More in detail, VERSO STEP #1 accounts for uncertainty in the data, by employing a maximum likelihood approach (via MCMC search) that allows for the presence of false positives, false negatives and missing data points in clonal variant profiles. As shown in^105^ in a different experimental context, our algorithmic framework ensures robustness and scalability also in case of high rates of errors and missing data, due for instance to sampling limitations. Furthermore, it is robust to mild violations of the ISA, e.g., due to reticulation events (e.g., convergent variants) or mutation losses, which can be characterized after the inference, if present (see the specific features on homoplasy detection in the next subsections). Please refer to the Supplementary Material for further details on the algorithmic framework and its assumptions, including the probabilistic graphical model depicted in Supplementary Figure S1 and the summary of notation in Table S4 of the Supplementary Material.

#### Outputs

The inference returns a set of maximum likelihood variants trees (minimum 1) as sampled during the MCMC search, representing the ordering of accumulation of clonal variants, and a set of maximum likelihood attachments of samples to variants. Given the variants tree and the maximum likelihood attachments of samples to variants, VERSO outputs: (*i*) a phylogenetic model where each leaf correspond to a sample, whereas internal nodes correspond to accumulating clonal variants, (*ii*) the corrected clonal genotype of each sample, i.e., the binary mutational profile on clonal variants obtained after removing false positives, false negatives and missing data.

The model naturally includes polytomies, which group samples with the same corrected clonal genotype. The length of the branches in the model represents the number of clonal substitutions (which can be normalized with respect to genome length), as in standard phylogenomic models, and the clades of the model correspond to viral lineages. The VERSO phylogenetic model is provided as output in Newick file format and can be processed and visualized in standard tools for phylogenetic analysis, such as FigTree^104^ or Dendroscope^108^. Furthermore, VERSO allows one to visualize the geo-temporal localization of clonal genotypes via Microreact^87^.

#### Additional feature: homoplasy detection on clonal variants

Violations of the ISA are possible and can be due to reticulation events^53^ such as homoplasies, i.e., identical variants detected in samples belonging to different clades, or to rare occurrences involving mutation losses (e.g., due to recombination-related deletions or to multiple mutations hitting an already mutated genome location^34^), as well as to infrequent transmission phenomena, such as super-infections^65,66^ (for a discussion on the general limitations of approaches based on phylogenetic trees when dealing with reticulation events refer, e.g., to^109–111^).

In this regard, VERSO allows one to identify clonal mutations likely involved in homoplasies, in a similar fashion to the plethora of works on mitochondrial evolution and phylogenetic networks (see, for instance, ^53,54,56,112–114^). In detail, given the maximum likelihood phylogenetc tree, VERSO can estimate the variants that are theoretically expected in each sample. By comparing the theoretical observations with the input data, VERSO can estimate the rate of false positives (i.e., the variants that are observed in the data but are not predicted by VERSO), and false negatives (i.e., variants that are not observed, but predicted). Variants that show a particularly high level of estimated error rates represent candidate homoplasies and are flagged. First, this allows one to pinpoint samples exhibiting homoplastic mutations (see Figure 3 and Supplementary Figures S8) and, second, to reconstruct an *expanded* clonal variants tree, in which candidate homoplastic mutations are duplicated after the inference, so to allow the visualization of reticulation events, as proposed in ^112^ (see, e.g., Supplementary Figures S3 and S9).

Furthermore, once this procedure has been completed, the list of flagged variants can include: (*i*) mutations falling in highly-mutated regions due to mutational hotspots, (*ii*) phantom mutations i.e., systematic artifacts generated during sequencing processes^56^, or (*iii*) mutations that have been positively selected in the population, e.g., due to a particular functional advantage.

Since one might be interested in identifying positively selected mutations, VERSO allows one to perform a consecutive analysis, which aims at highlighting the mutation-prone regions of the genome, and which might be due to mutational hotspots or phantom mutations (see the Supplementary Material for further details). We finally note that the detection of homoplasies for minor variants require a different algorithmic procedure, which is detailed in the following.

### VERSO STEP #2: Characterization of intra-host genomic diversity

In the second step, VERSO takes into account the variant frequency profiles of groups of samples with the same corrected clonal genotype (identified via VERSO STEP #1), in order to characterize their intra-host genomic diversity and visualize it on a low-dimensional space. This allows one to highlight patterns of co-occurrence of minor variants, possibly underlying undetected infection events, as well as homoplasies involving, e.g., positively selected variants. Notice that this step requires raw sequencing data and the prior execution of STEP #1.

#### Inputs

VERSO STEP #2 takes as input a *n* (samples) ×*m* (variants) variant frequency (VF) profile matrix, in which each entry includes the VF ∈ (0, 1) of a given mutation in a certain sample, after filtering out: (*i*) the clonal variants employed in STEP #1 and (*ii*) the minor variants possibly involved in homoplasies (see below). The variant calling pipeline employed in this work is detailed in the next subsections.

#### The algorithmic framework

While it is sound to binarize clonal variant profiles to reconstruct a phylogenetic tree, it is opportune to consider the variant frequency profiles when analyzing intra-host variants, for several reasons. First, variant frequency profiles describe the intra-host genomic diversity of any given host, and this information would be lost during binarization. Second, minor variant profiles might be noisy, due to the relatively low abundance and to the technical limitations of sequencing experiments. Accordingly, such data may possibly include artifacts, which can be partially mitigated during the quality-check phase and by including in the analysis only highly-confident variants. However, binarization with arbitrary thresholds might increase the false positive rate, compromising the accuracy of any downstream analysis. Third, as specified above, the extent of transmission of minor variants among individuals is still partially obscure. The VF of minor variants is, in fact, highly affected by recombination processes, as well as by complex transmission phenomena, involving stochastic fluctuations, bottlenecks and founder effects, and which may lead certain variants changing their VF, not being transmitted or even becoming clonal in the infected host^57^. The latter issue also suggests that the hypothesis of accumulation of minor variants during infections may not hold and should be relaxed.

For these reasons, VERSO STEP #2 defines a pairwise genomic distance, computed on the variant frequency profiles, to be used in downstream analyses. The intuition is that samples displaying similar patterns of co-occurrence of minor variants might have a similar quasispecies architecture, thus being at a small evolutionary distance. Accordingly, this might indicate a direct or indirect infection event. In particular, in this work we employed the Bray-Curtis dissimilarity, which is defined as follows: given the ordered VF vectors of two samples, i.e. 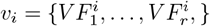 and 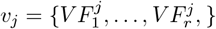, the pairwise Bray-Curtis dissimilarity *d*(*i, j*) is given by:

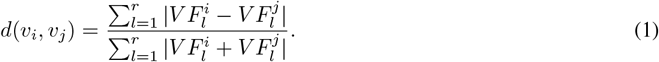

Since this measure weights the pairwise VF dissimilarity on each variant with respect to the sum of the VF of all variants detected in both samples, it can be effectively used to compare the intra-host genomic diversity of samples, as proposed for instance in^115^. However, VERSO allows one to employ different distance metrics on VF profiles, such as correlation or Euclidean distance.

As a design choice, in VERSO the genomic distance is computed among all samples associated to any given corrected clonal genotype, as inferred in STEP #1. The rationale is that, in a statistical inference framework modeling a complex interplay involving heterogeneous dynamical processes, it is crucial to stratify samples into homogeneous groups, to reduce the impact of possible confounding effects^116^. Furthermore, as specified above, due to the distinct properties of clonal and minor variants during transmission, it is reasonable to assume that the event in which certain minor variants and no clonal variants are transmitted from a host to another during the infection is extremely unlikely. Accordingly, the clonal variants employed for the reconstruction of the phylogenetic tree in STEP #1 are excluded from the computation of the intra-host genomic distance among samples.

In order to produce useful knowledge from the genomic distance discussed above and since, in real-world scenarios, this is a typically complex high-dimensional problem, it is sound to employ state-of-the-art strategies for dimensionality reduction and (sample) clustering, as typically done in single-cell analyses^117^. In this regard, the workflow employed in VERSO ensures high scalability with large datasets, also allowing to taking advantage of effective analysis and visualization features. In detail, the workflow includes three steps: (*i*) the computation of the *k-nearest neighbour* graph (k-NNG), which can be executed on the original variant frequency matrix, or after applying *principal component analysis* (PCA), to possibly reduce the effect of noisy observations (when the number of samples and variants is sufficiently high); (*ii*) the clustering of samples via either *Louvain* or *Leiden* algorithms for community detection^118^; (*iii*) the projection of samples on a low-dimensional space via standard tSNE^68^ or UMAP^67^ plots.

#### Outputs

As output, VERSO STEP #2 delivers both the partitioning of samples in homogeneous clusters and the visualization in a low-dimensional space, also allowing to label samples according to other covariates, such as, e.g., collection date or geographical location. In the map in Fig. 3, for instance, the intra-host genomic diversity of each sample and the genomic distance among samples are projected on the first two UMAP components, whereas samples that are connected by k-NNG edges display similar patterns of co-occurrence of variants. Accordingly, the map shows clusters of samples likely affected by infection events, in which (a fraction of) quasispecies might have been transmitted from a host to another. This represents a major novelty introduced by VERSO and also allows one to effectively visualize the space of variant frequency profiles.

To facilitate the usage, VERSO STEP #2 is provided as a Python script which employs the SCANPY suite of tools^59^, which is typically used in single-cell analyses and includes a number of highly-effective analysis and visualization features.

#### Additional feature: homoplasy detection on minor variants

Also in the case of minor variants, it is important to pinpoint possible homoplasies and which might be due to mutational hotspots, phantom mutations and convergent variants. Given the phylogenetic model retrieved via STEP #1, VERSO allows one to flag the variants that are detected in a number of clonal genotypes exceeding a user-defined threshold. In our case, the threshold is equal to 1, meaning that all minor variants found in more than one clonal genotypes are flagged.

Such variants are then excluded from the computation of the intra-host genomic distance, prior to the execution of STEP #2. Furthermore, the list of flagged variants can be investigated as proposed for STEP #1 (see above), in order to possibly identify mutations involved in positive selection scenarios.

## Datasets description

### Dataset #1 (Illumina Amplicon sequencing)

We analyzed 3960 samples from distinct individuals obtained from 22 NCBI BioProjects, which, at the time of writing, are all the publicly available datasets including raw Illumina Amplicon sequencing data. In detail, we selected the following projects: (1) PRJNA613958, (2) PRJNA614546, (3) PRJNA616147, (4) PRJNA622817, (5) PRJNA623683, (6) PRJNA625551, (7) PRJNA627229, (8) PRJNA627662, (9) PRJNA629891, (10) PRJNA631042, (11) PRJNA633948, (12) PRJNA634119, (13) PRJNA636446, (14) PRJNA636748, (15) PRJNA639066, (16) PRJNA643575, (17) PRJNA645906, (18) PRJNA647448, (19) PRJNA647529, (20) PRJNA650037, (21) PRJNA656534, (22) PRJNA656695.

### Dataset #2 (Illumina RNA-sequencing)

We analyzed 2766 samples from distinct individuals obtained from 22 NCBI BioProjects, which, at the time of writing, are all the publicly available datasets including raw Illumina RNA-sequencing data. In detail, we selected the following projects: (1) PRJNA601736, (2) PRJNA603194, (3) PRJNA605983, (4) PRJNA607948, (5) PRJNA608651, (6) PRJNA610428, (7) PRJNA615319, (8) PRJNA616446, (9) PRJNA623895, (10) PRJNA624792, (11) PRJNA626526, (12) PRJNA631061, (13) PRJNA636446, (14) PRJNA637892, (15) PRJNA639591, (16) PRJNA639864, (17) PRJNA650134, (18) PRJNA650245, (19) PRJNA655577, (20) PRJNA657938, (21) PRJNA657985, (22) PRJNA658211.

### Contact tracing data

Contact tracing data were obtained from the study presented in^69^. In detail, for 65 samples included in Dataset #1 (NCBI BioProject PRJNA633948), information on households, work institutions and epidemiological linkages are provided. Thus, it is possible to identify 3 different contact groups based on institutions regularly frequented by patients and 1 household couples. Contact information was employed to assess the relation between the intra-host genomic similarity and the contact dynamics. The results are provided in the main text.

## Parameter settings

### Parameter settings of variant calling (Datasets #1 and #2)

We converted all the samples to FASTQ files using SRA toolkit. Following^75^, we used Trimmomatic (version 0.39) to remove the nucleotides with low quality score from the RNA sequences with the following settings: LEADING:20 TRAILING:20 SLIDINGWINDOW:4:20 MINLEN:40. We then used bwa mem (version 0.7.17) to map reads to SARS-CoV-2-ANC reference genome (Supplementary File S1; see Results). We generated sorted BAM files from bwa mem results using SAMtools (version 1.10) and removed duplicates with Picard (version 2.22.2). Variant calling was performed generating mpileup files using SAMtools and then running VarScan (min-var-freq parameter set to 0.01)^119^.

We note that it was recently reported that some currently available SARS-CoV-2 datasets exhibit quality issues^13,120^. Accordingly, one should be extremely careful when performing quality check and, especially, when considering low-frequency variants, which might possibly result from sequencing artifacts even in case of high-coverage experiments. In this regard, many effective approaches can be employed to reduce false variants. For instance, the Broad Institute recently updated an effective variant calling pipeline for viral genome data^121^, while new methods for error correction of viral sequencing have been proposed at this widely used website: https://virological.org, which also includes a number of useful up-to-date guidelines and best practices for viral evolution analyses.

In our case, we here employed the following significance filters on variants. In particular, we kept only the mutations: (1) showing a VarScan significance p-value *<* 0.01 (Fisher’s Exact Test on the read counts supporting reference and variant alleles) and more than 25 reads of support in at least 75% of the samples, (2) displaying a variant frequency VF > 5%. As a result, we selected a list of 15892 (over 55280 overall SNVs) highly-confident SNVs for Dataset #1 and 7389 (over 53354) for Dataset #2.

High-quality variants were then mapped on SARS-CoV-2 coding sequences (CDSs) via a custom R script, also by highlighting synonymous/nonsynonymous states and amino acid substitutions for the related Open Reading Frame (ORF) product. In particular, we translated reference and mutated CDSs with the seqinr R package to obtain the relative amino acid sequences, which we compared to assess the effect of each nucleotide variation in terms of amino acid substitution.

We finally note that availability of the Ct values generated by Q-PCR and the related quantification of the amount of viral transcripts would be very useful to characterize samples with high viral load, yet this information is not available for the considered datasets.

### Quality check (Datasets #1 and #2)

In order to select high-quality samples, we selected only those exhibiting high coverage and in particular those with at least 25 reads in more than 90% of the SARS-CoV-2-ANC genome. In addition, we filtered out all samples exhibiting more than 100 minor variants (VF ≤ 90%).

We finally excluded samples SRR11597146 and SRR11476447 from Dataset #1, as the first sample displays zero SNVs and the second one reports an unfeasible collection date (i.e. 30^th^ Jan. 2019).

After the quality-check filters, 2906 samples of Dataset #1 are left for downstream analyses, in which 10571 distinct high-quality single-nucleotide variants are observed, and 1438 samples are left for Dataset #2, with 6143 high-quality SNVs.

### Parameter settings of VERSO (Datasets #1 and #2)

The phylogenomic analysis via VERSO STEP #1 was performed on Datasets #1 and #2 by considering only clonal variants (VF > 90%) detected in at least 3% of the samples. A grid search comprising 16 different error rates was employed (see Table S3 of the Supplementary Material). Samples with the same corrected clonal genotype were grouped in polytomies in the final phylogenetic models.

The analysis of the intra-host genomic diversity via VERSO STEP #2 was performed by considering the VF profiles of all samples, by excluding: (*i*) the clonal variants employed in the phylogenomic reconstruction via VERSO STEP #1, (*ii*) the minor variants involved in homoplasies, i.e., observed in more than one clonal genotype returned by VERSO STEP #1. Missing values (NA) were imputed to 0 for downstream analysis. A number of PCs equals to 10 was employed in PCA step, prior to the computation of the k-nearest neighbour graph (*k* = 10) on the Bray-Curtis dissimilarity of VF profiles. Leiden algorithm was applied with resolution = 1 (see Table S3 of the Supplementary Material for the parameter settings of VERSO employed in the case studies).

### Parameter settings of simulations

In order to compare the performance of VERSO STEP #1 with competing phylogenomic tools, i.e., IQ-TREE^10^ and BEAST 2^22^, we performed extensive simulations via msprime^70^, which simulates a backwards-in-time coalescent model.

In particular, we simulated 20 distinct evolutionary processes, with the following parameters: *n* = 1000 total samples, effective population size *N*_*e*_ = 0.5 (i.e., haploid population), mutational rate *M* = 2 ×10^−6^ mutations per site per generation and a genome of length *L* = 29903 bases. Such parameters were chosen to roughly approximate the mutational rate currently estimated for SARS-CoV-2 (i.e., *M* ≈10^*−*^3 mutations per site per year and 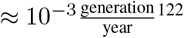) and to obtain a number of clonal mutations (in the range 15 − 30) that is comparable to the one observed in the real-word scenarios (see the case studies). As output, msprime returns a phylogenetic tree representing the genealogy between the samples, the genotype of all samples (i.e., the leaves of the tree) and the location of all mutations.

The genotypes of the samples were then inflated with different levels of noise, with false positive rate *α* and false negative rate *β* (see the parameter settings in Table S1 of the Supplementary Material), in order to assess the performance of the methods in conditions of noisy observations and possible sequencing issues. Finally, we subsampled all datasets to obtain two distinct samples sizes (500 and 1000 samples), in order to test the robustness of methods in conditions of sampling limitations.

The parameters of the phylogenetic methods employed in the comparative assessment are reported in the Supplementary Material (Table S2 of the Supplementary Material).

## Supporting information

Graphical Abstract

## Acknowledgments

This work was partially supported by the Elixir Italian Chapter and the SysBioNet project, a Ministero dell’Istruzione, dell’Università e della Ricerca initiative for the Italian Roadmap of European Strategy Forum on Research Infrastructures and by the AIRC-IG grant 22082. Partial support was also provided by the CRUK/AECC/AIRC Accelerator Award #22790, “Single-cell Cancer Evolution in the Clinic”. We thank Giulio Caravagna, Chiara Damiani, Lucrezia Patruno and Francesco Craighero for helpful discussions. We also thank David Posada for interesting suggestions on the preliminary version of the manuscript.

## Authors contributions

D.R., F.A., D.M., A.G. and R.P. designed the approach. D.R., F.A., D.M. and A.G. defined, implemented and executed the computational methods. D.R., F.A., D.M. and A.G. performed the simulations. D.R., F.A., D.M., C.G., M.A., A.G. and R.P. analyzed the data and interpreted the results. R.P. supervised the experimental data analysis. A.G. and D.R. supervised the computational analysis. A.G. and R.P. drafted the manuscript, which all authors discussed, reviewed and approved.

## Supplementary Files Captions

- **Supplementary File S**1. SARS-CoV-2-ANC reference genome in FASTA format.
- **Supplementary File S**2. Summary of the samples exhibiting homoplastic clonal variants in Datasets #1 and #2.
- **Supplementary File S**3. Mapping between the clonal genotype labels of the phylogenomic models returned via VERSO STEP #1 on Datasets #1 and #2 and the lineage dynamic nomenclature proposed in^78^, generated via pangolin 2.0^79^.

