## Supplementary figures and images for "VERSO: a comprehensive framework for the inference of robust phylogenies and the quantification of intra-host genomic diversity of viral samples"

### Graphical Abstract

#### V E R S O | Viral Evolution ReconStructiOn

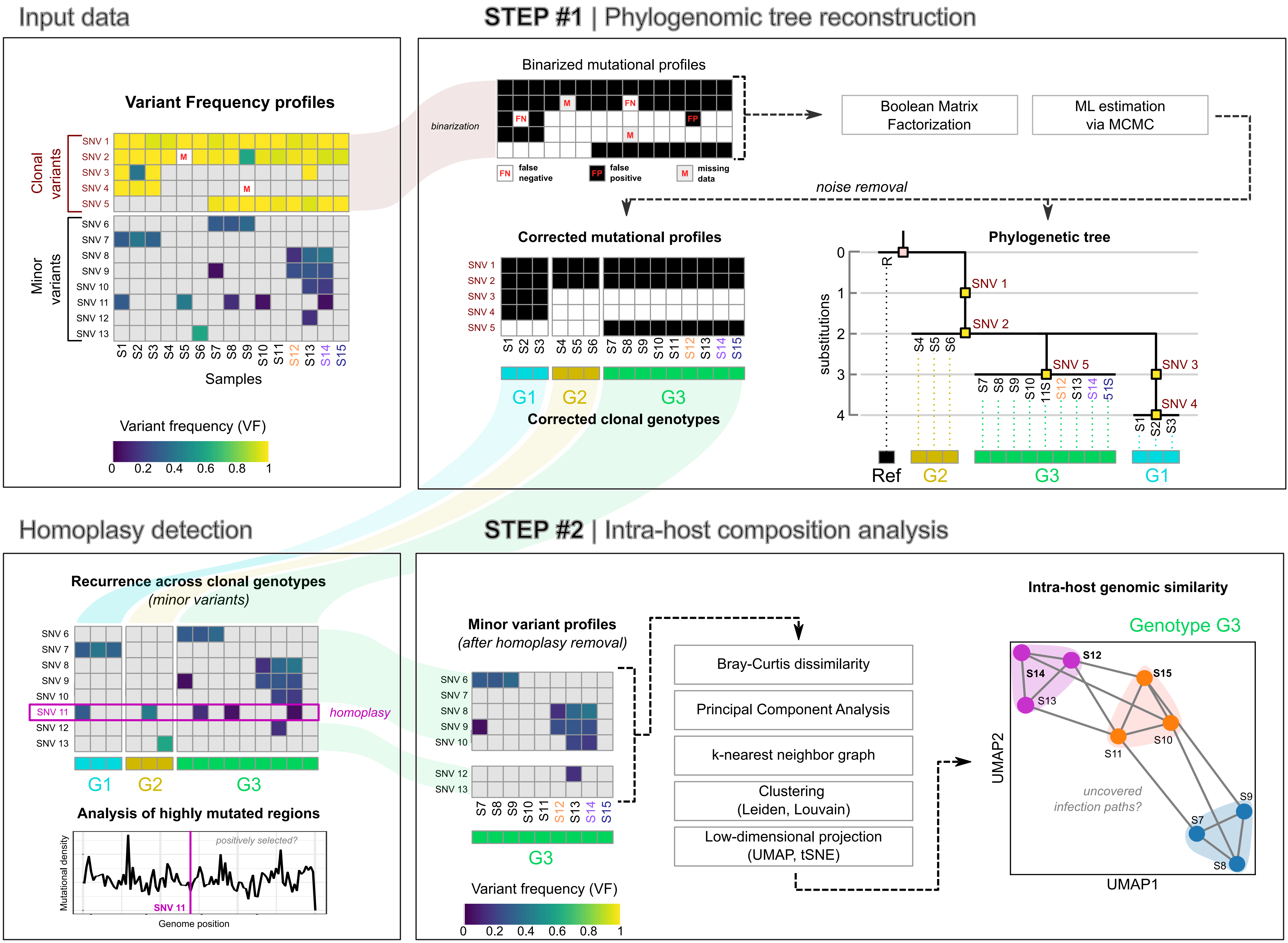
