## Supplementary Information for "VERSO: a comprehensive framework for the inference of robust phylogenies and the quantification of intra-host genomic diversity of viral samples"

### Contents


**1 Additional materials and methods**

1.1 Additional details on VERSO STEP #1 algorithmic framework

1.2 Performance assessment metrics

1.3 Parameter settings

1.4 Summary of notation

**2 Additional results**

2.1 Reference genome generation

2.2 Expanded clonal variants tree – Dataset #1

2.3 Mutational hotspot analysis – Dataset #1

2.4 Bottleneck size estimation – Dataset #1

2.5 Impact of clonal variant threshold on VERSO stability – Dataset #1

2.6 Application of VERSO to 1438 samples from RNA sequencing data (Dataset #2)

2.7 Expanded clonal variants tree – Dataset #2

2.8 Computation time


### 1 Additional materials and methods

#### 1.1 Additional details on VERSO STEP #1 algorithmic framework

VERSO STEP #1 aims at inferring robust phylogenomic models from either raw sequencing data or consensus sequences, and employs an algorithmic strategy akin to[1]. In particular, VERSO STEP #1 takes as input either: $(i)$ (if raw sequencing data are available) the binarized profiles of clonal SNVs detected in the cohort, which are selected via a variant frequency (VF) threshold (equal to 90% in the case studies) or, $(ii)$ the consensus sequences of the samples in the cohort, which are translated in opportune binarized clonal variants profiles by employing a reference genome (see the main text).

More in detail, let us consider $k$ different clonal genotypes characterized by $m$ clonal variants and $n$ samples $s_1, \ldots, s_n$. We define the *theoretical clonal genotype matrix* $\mathbf{G}$: a binary $n \times m$ matrix where each row represents a sample and each column a clonal mutation; an element of $\mathbf{G}$, $g_{i,j} = 1$ if variant $j$ is present in sample $i$, otherwise $g_{i,j} = 0$. We then define the *phylogenetic matrix* $\mathbf{B}$: a binary $k \times m$ matrix where each row represents a clonal genotype and each column a clonal mutation; $b_{ij} = 1$ if we detect mutation $j$ in clonal genotype $i$, otherwise $b_{i,j} = 0$. Each $\mathbf{B}$ uniquely identifies a clonal variants tree[2]. We here assume the existence of a perfect phylogeny process and that the Infinite Sites Assumption holds[3], i.e., mutations cannot be lost, they can compare only once in the tree, and the tree has only one root. Notice that violations of the Infinite Sites Assumption are possible due to, e.g., homoplasies or loss of mutations, which are characterized after the inference (see the specific features on homoplasy detection described in the main text).

Finally, we define the *sample attachment matrix* $\mathbf{S}$: a binary $n \times k$ matrix, where each row represents a sample and each column a clonal genotype; $s_{i,j} = 1$ if sample $i$ is associate to clonal genotype $j$, otherwise $s_{i,j} = 0$; each sample is attached exactly to one clonal genotype.

Therefore, the following factorization holds:

$$\mathbf{G} = \mathbf{S} \cdot \mathbf{B}. \tag{1}$$

In general, the observed clonal genotype of a given sample $i$ in real-world data may not correspond to its theoretical clonal genotype (the $i$-th row of $\mathbf{G}$), because it can include include false positives, false negatives and missing values. Therefore, when considering a real-word data matrix ($\mathbf{D}$), the previous equation typically does not hold. For this reason, VERSO employs a maximum likelihood estimation procedure to determine the model that best fits the data, given an appropriate noise model. Let us define the data matrix $\mathbf{D}$ a binary $n \times m$ matrix where each row represents a sample and each column a clonal mutation; an element of $\mathbf{D}$, $d_{i,j} = 1$ if the variant $j$ is present in sample $i$, otherwise $d_{i,j} = 0$. Then, the posterior probability of an output model (i.e., $\mathbf{B}$, $\mathbf{S}$), and of error rates $\alpha$ (i.e., the false positive rate) and $\beta$

(i.e., the false negative rate), given the data matrix $\mathbf{D}$ and the hyperparameters of error rate distributions $\theta_\alpha$ and $\theta_\beta$ can be written as:

$$P(\mathbf{B},\mathbf{S},\mathbf{G},\alpha,\beta|\mathbf{D},\theta_\alpha,\theta_\beta) \propto P(\mathbf{D}|\mathbf{B},\mathbf{S},\mathbf{G},\alpha,\beta,\theta_\alpha,\theta_\beta)P(\mathbf{B},\mathbf{S},\mathbf{G},\alpha,\beta|\theta_\alpha,\theta_\beta). \quad (2)$$

We then assume independence of both the phylogenetic model $\mathbf{B}$ and the sample attachment $\mathbf{S}$ from error rates and a uniform uninformative prior on $\theta_\alpha$ and $\theta_\beta$. Furthermore, we assume that the data matrix $\mathbf{D}$ depends only on theoretical clonal genotype matrix $\mathbf{G}$ and error rates $\alpha$ and $\beta$, we obtain:

$$P(\mathbf{B},\mathbf{S},\mathbf{G},\alpha,\beta|\mathbf{D},\theta_\alpha,\theta_\beta) \propto P(\mathbf{D}|\mathbf{G},\alpha,\beta)P(\mathbf{G}|\mathbf{B},\mathbf{S})P(\mathbf{S}|\mathbf{B})P(\mathbf{B})P(\alpha|\theta_\alpha)P(\beta|\theta_\beta). \quad (3)$$

Finally, assuming: *i)* an uniform uninformative prior on $\mathbf{B}$, *ii)* that given the hyper-parameters of error rates, all possible values of $\alpha$ and $\beta$ are equiprobable (this because, in the current formulation, VERSO includes a noise estimation procedure via grid search over a limited number of values of $\alpha$ and $\beta$), and *iii)* that given any $\mathbf{B}$, all $\mathbf{S}$ are equiprobable, we obtain:

$$P(\mathbf{B},\mathbf{S},\mathbf{G},\alpha,\beta|\mathbf{D},\theta_\alpha,\theta_\beta) \propto P(\mathbf{D}|\mathbf{G},\alpha,\beta). \quad (4)$$

Then, the problem is reduced to the maximization of the following likelihood function:

$$P(\mathbf{B},\mathbf{S},\mathbf{G},\alpha,\beta|\mathbf{D},\theta_\alpha,\theta_\beta) \propto \prod_{i=1}^{n}\prod_{j=1}^{m} P(d_{i,j}|\mathbf{G}_{i,j},\alpha,\beta), \quad (5)$$

where $d_{i,j}$ is a entry of $\mathbf{D}$ and:

$$P(d_{i,j}|\mathbf{G}_{i,j},\alpha,\beta) = \begin{cases} \alpha, & if \quad d_{i,j}=1 \quad and \quad \mathbf{G}_{i,j}=0, \\ 1-\alpha, & if \quad d_{i,j}=1 \quad and \quad \mathbf{G}_{i,j}=1, \\ \beta, & if \quad d_{i,j}=0 \quad and \quad \mathbf{G}_{i,j}=1, \\ 1-\beta, & if \quad d_{i,j}=0 \quad and \quad \mathbf{G}_{i,j}=0, \\ 1, & if \quad d_{i,j}=\text{NA}, \end{cases} \quad (6)$$

where NA label missing entries, e.g., due to low coverage. In Figure S1 the graphical probabilistic model of VERSO is shown .

As an exhaustive search can be performed only for very small models, we employ a Markov Chain Monte Carlo (MCMC) scheme via a Metropolis–Hastings algorithm, to search the model that maximizes the likelihood function defined in (5) . The MCMC includes two ergodic moves on $\mathbf{B}$ : $(i)$ node relabeling, $(ii)$ prune/reattach of a single node and its descendants. For each proposed configuration $\mathbf{B}'$, we find the maximum likelihood $\mathbf{S}'$ via an exhaustive search and we then evaluate the new proposed corrected clonal genotype $\mathbf{G}'$. Therefore, the acceptance ratio of a move $\rho_{\mathbf{B}',\mathbf{S}',\mathbf{G},\alpha,\beta}$ is given by:

$$\rho_{\mathbf{B}',\mathbf{S}',\mathbf{G},\alpha,\beta} = \min\left\{ \left(\frac{P(\mathbf{D}|\mathbf{G}',\alpha,\beta)}{P(\mathbf{D}|\mathbf{G},\alpha,\beta)}\right)^{1/T}, 1\right\}, \quad (7)$$

where $T$ is a learning rate parameter, which can be tuned as suggested in[4] (default $T=1$).

Once the maximum likelihood model ($\mathbf{B}$, $\mathbf{S}$, $\alpha$ and $\beta$) is found, the clonal variants tree $\mathbf{B}$ represents the inner structure of the output phylogenetic tree in which variants are the inner nodes, whereas the sample attachment matrix $\mathbf{S}$ determines the edges connecting the inner nodes to the samples, which are the leaves of the phylogenetic tree. Since the number of corrected clonal genotypes is typically much lower than the number of samples, samples with the same corrected genotype are grouped in polytomies, i.e., they are children of the same inner node.

Notice that, in the output phylogenetic tree, the branch length correspond to the number of clonal substititions, which can be normalized according to the genome length. Finally, in addition to the maximum likelihood phylogenetic model, VERSO also provides as output the corrected clonal genotype of all samples.

##### 1.2 Performance assessment metrics

The performance of the different methods was assessed by simulations, comparing the inferred phylogenies with the respective ground-truth topology; three metrics were considered (i) absolute error evolutionary distance, (ii) branch score difference[5] and (iii) quadratic path difference[6], which are detailed in the following.

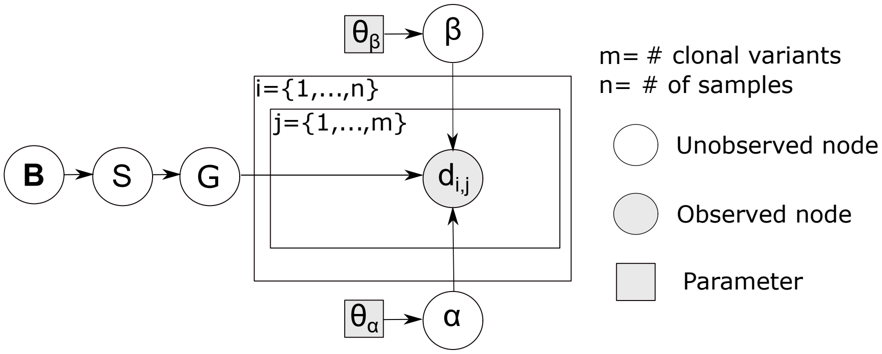


Figure S1: **Probabilistic graphical model of VERSO STEP #1.** Shaded nodes represent observed values. Square and shaded nodes represent fixed parameters; unshaded nodes represent latent variables. The probability distribution of unshaded nodes is approximated using samples from the proposed search scheme. Please refer to the text for the definition of variables and parameters.

$(i)$ **Absolute Error Evolutionary Distance.** For two phylogenetic trees, we compute the pairwise distances between each pairs of tips using its branch lengths. The Absolute Error Evolutionary Distance is the mean absolute error between such distances in the two trees.

$(ii)$ **Branch Score Difference.** To compute this distance, we consider the list of all possible partitions over two phylogenetic trees, some of which will be found in the trees, while others correspond to branches that are not admissible in some of them; branch lengths of 0 are assigned to these latter partitions. For the two trees, the Branch Score Difference[5] is the sum of squared differences between such branch lengths.

$(iii)$ **Quadratic Path Difference.** The length of a path connecting one tip to another in a phylogenetic tree is the number of edges that must be navigated to get from the first tip to the other. Given two phylogenetic trees, the Quadratic Path Difference[6] is the square root of the sum of squares of the difference in path length between each pair of tips.

##### 1.3 Parameter settings

| **Parameters of simulations via msprime[7]** | | | | |
|---|---|---|---|---|
| **# of simulations** | 20 | | | |
| **Sample size n** | 1000 | | | |
| **Effective population size** | 0.5 | | | |
| **Mutational rate** | $2 \times 10^{-6}$ mutations per generation | | | |
| **Genome length** | 29903 | | | |
| **Features of the sythetic datasets** | | | | |
| **Scenario** | A | B | C | D |
| **# of datasets** | 20 | 20 | 20 | 20 |
| **# of Samples** | 1000 | 1000 | 500 | 500 |
| false positive rate $\alpha$ | 0.05 | 0.1 | 0.05 | 0.1 |
| false negative rate $\beta$ | 0.05 | 0.1 | 0.05 | 0.1 |

Table S1: Parameter settings of the coalescent simulations via msprime[7] and of the related datasets employed in the comparative assessment presented in the main text.

| Settings of phylogenetic methods on simulated datasets | |
|---|---|
| **VERSO STEP $\#1$** | |
| **MCMC iterations** | 100000 |
| **# of MCMC restarts** | 500 |
| **Early stop threshold** | 5000 |
| **Learning rate $T$** | 1 |
| **Grid search $\alpha$** | $[0.001, 0.010, 0.050, 0.100]$ |
| **Grid search $\beta$** | $[0.001, 0.010, 0.050, 0.100]$ |
| **IQ-TREE** - Launched via Nextstrain-Augur pipeline | |
| **Software version** | 1.6.12 |
| **Substitution model** | GTR |
| **Other parameters** | Default |
| **BEAST 2** | |
| **Software version** | 2.6.3 |
| **MCMC iterations** | $1 \times 10^6$ |
| **Substitution model** | GTR |
| **Clock rate** | fixed (=1) |
| **Population model** | costant (=1) |
| **All priors** | default |
| **All move probability** | default |
| **Burn-in of trees** | 10% |
| **Target tree type** | maximun clade credibility |
| **Node heights** | median |

Table S2: Parameters settings of VERSO STEP $\#1$, IQ-TREE and Beast 2 with respect to the comparative assessment on simulations presented in the main text. IQ-TREE was launched via Nextstrain-Augur pipeline[8].

| **Features of real-world datasets** | | |
|---|---|---|
| **Dataset** | Dataset #1 | Dataset #2 |
| **Sequencing protocol** | Amplicon | RNA-seq |
| **# of samples - pre QC** | 3960 | 2766 |
| **# of samples - post QC** | 2906 | 1438 |
| **# of variants - pre QC** | 55280 | 53354 |
| **# of variants - post QC** | 10571 | 6143 |
| **# of clonal variants - included in the inference** | 29 | 23 |
| **Clonal threshold $\delta$** | 0.9 | 0.9 |
| **VERSO STEP #1: parameter settings** | | |
| **MCMC iterations** | 100000 | 100000 |
| **# of MCMC restarts** | 500 | 500 |
| **Early stop threshold** | 5000 | 5000 |
| **Learning rate $T$** | 1 | 1 |
| **Grid search $\alpha$** | $[0.001, 0.010, 0.050, 0.100]$ | $[0.001, 0.010, 0.050, 0.100]$ |
| **Grid search $\beta$** | $[0.001, 0.010, 0.050, 0.100]$ | $[0.001, 0.010, 0.050, 0.100]$ |
| **VERSO STEP #2: parameter settings** | | |
| **Distance type** | Bray-Curtis diss. | Bray-Curtis diss. |
| **# of PCAs** | 10 | 10 |
| **# of $k$ (kNN)** | 10 | 10 |
| **Leiden resolution** | 1 | 1 |
| **Spread (UMAP)** | 1 | 1 |
| **Min_dist (UMAP)** | 2 | 2 |
| **a (UMAP)** | 0.01 | 0.01 |
| **b (UMAP)** | 1 | 1 |

Table S3: Features of the real-world datasets employed in the case studies and parameter settings of both steps of VERSO with respect to the application of such datasets.

##### 1.4 Summary of notation

| Symbol | Description |
|---|---|
| $m$ | Number of clonal variants |
| $k$ | Number of clonal genotypes in the model |
| $n$ | Number of samples |
| $\alpha$ | False Positive Rate |
| $\beta$ | False Negative Rate |
| $\gamma$ | Ratio of missing entries |
| $\delta$ | Clonal variant VF threshold |
| $\mathbf{G}$ | Theoretical clonal genotype matrix |
| $\mathbf{B}$ | Perfect phylogeny matrix |
| $\mathbf{D}_i$ | Input binary clonal variant profile matrix |
| $\mathbf{S}$ | Sample attachment matrix |
| $b_{i,j}$ | $i, j$ element of the matrix $\mathbf{B}$ |
| $d_{i,j}$ | $i, j$ element of the matrix $\mathbf{D}$ |
| $v_i$ | ordered vector of the VF of the $i$-th sample |
| $s_{i,j}$ | $i, j$ element of the matrix $\mathbf{S}$ |
| $\mathbf{b}_i$ | $i$-th row of the phylogenetic matrix $\mathbf{B}$ |
| $\rho_{\mathbf{B}',\mathbf{S}',\mathbf{G},\alpha,\beta}$ | Ratio of acceptance of a (i.e., the proposal of a $\mathbf{B}'$ and $\mathbf{C}'$ ) in the MCMC |
| $T$ | Learning rate parameter in the MCMC |
| $d(a, b)$ | Bray-Curtis dissimilarity between $a$ and $b$ |
| TP | True positive |
| FP | False positive |
| TN | True negative |
| FN | False negative |
| $\mathrm{VF}_i^j$ | Variant frequency of the $j$-th sample in the $i$-th nucleotide |

Table S4: Summary of the main notation used in the article.

### 2 Additional results

##### 2.1 Reference genome generation

The description of the generation of reference genome SARS-CoV-2-ANC is provided in the main text and depicted in Figure S2.

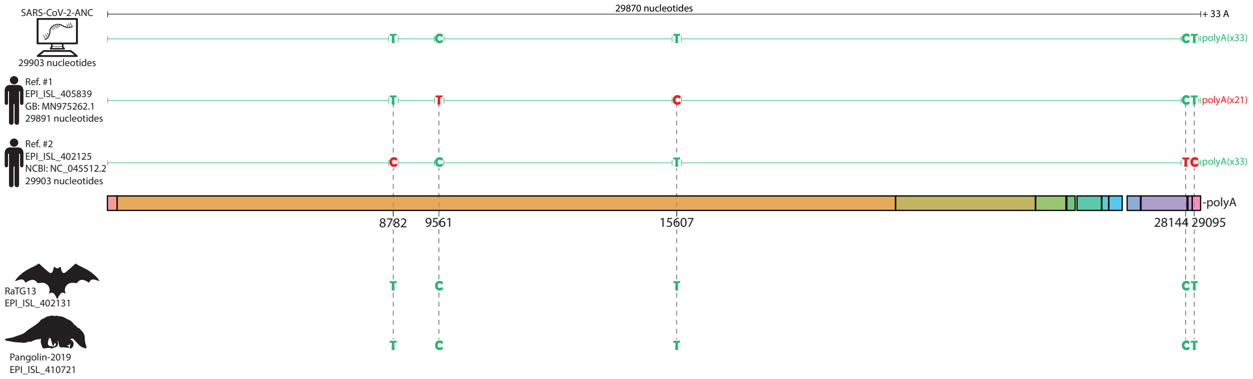


Figure S2: **Generation of SARS-CoV-2-ANC reference genome.** The artificial reference genome SARS-CoV-2-ANC was generated setting 29865 (on 29870) genome locations identical to both ref #1 (`EPI_ISL_405839`) and ref #2 (`EPI_ISL_402125`), including the polyA tail of ref. #2 (33 bases), and setting haplotype TCTCT at locations 8782, 9561, 15607, 28144 and 29095, which is shared by both the Bat-CoV-RaTG13 genome (sequence `EPI_ISL_402131`) and the Pangolin-CoV genome (sequence `EPI_ISL_410721`). The final length of the SARS-CoV-2-ANC genome is 29903.

##### 2.2 Expanded clonal variants tree – Dataset #1

In Figure S3 we report the *expanded* clonal variants tree of Dataset #1, obtained by duplicating homoplastic variants identified after the phylogenetic tree inference via VERSO STEP #1. In particular, the 5 homoplastic variants were duplicated in the tree, if observed in at least 10 samples with the same corrected clonal genotype, and were positioned after the corresponding characterizing mutation, as proposed in[9]. Homoplastic variants were finally connected via dashed lines to pinpoint reticulation events. In Figure 3 of the main text, all samples displaying homoplastic variants are highlighted directly on the phylogenetic model returned by VERSO STEP #1.

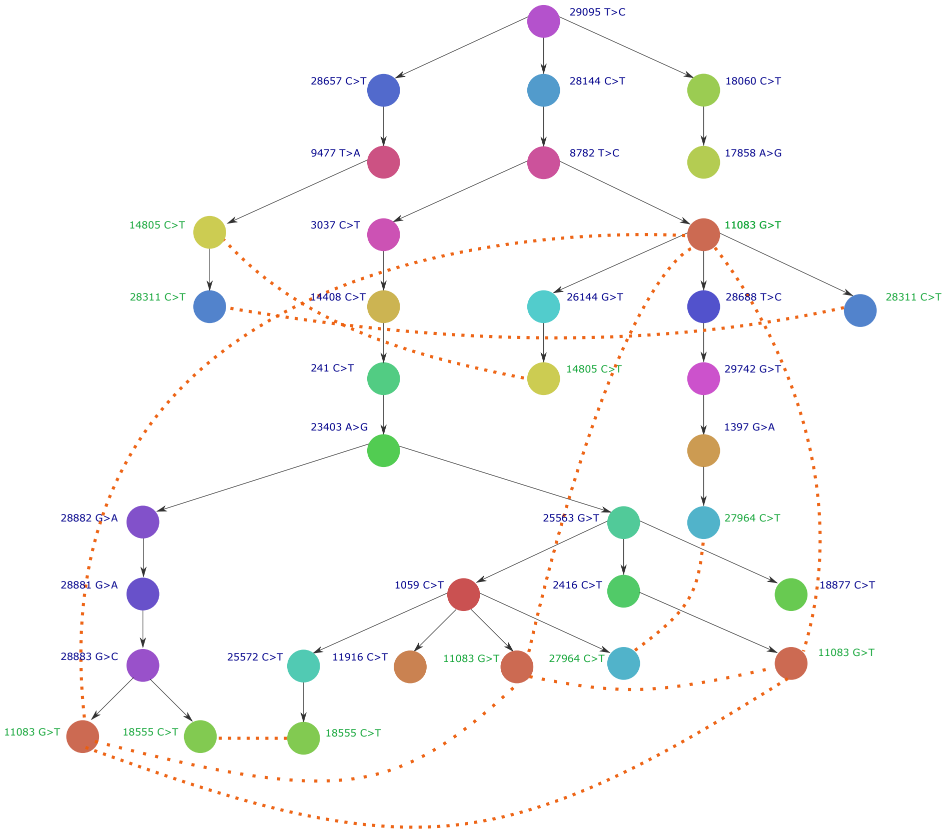


Figure S3: **Expanded clonal variants tree – Dataset #1.** The clonal variants tree **B** representing the inner structure of the output phylogenetic tree inferred from Dataset #1 via VERSO STEP #1 is expanded by duplicating homoplastic variants. Nodes represent clonal variants (consistent colors with Figure 3 of the Main Text), whereas edges indicate temporal ordering/accumulation relations. Homoplastic mutations were duplicated if observed in at least 10 samples with the same corrected clonal genotype returned by VERSO STEP #1 and are attached after the corresponding characterizing mutation. Homoplastic variants observed in multiple samples with the same corrected clonal genotype are ordered by the number of samples in which they are detected. Dashed lines connect identical homoplastic mutations.

##### 2.3 Mutational hotspot analysis – Dataset #1

To identify the regions of the SARS-CoV-2 genome more prone to mutations, which might be due to *natural* mutational hotspots or to *phantom mutations*, i.e., systematic artifacts generated during sequencing processes[10], it is sound to evaluate the mutation density with a sliding-window approach, as proposed, for instance, in[11]. This analysis is also effective to distinguish homoplasies due to positive selection from those due to artifacts or hotspots.

More in detail, the analysis considers: $(i)$ minor variants (i.e., with VF$< 90\%$) to prevent possible biases due to the transmission and accumulation of clonal variants in the population, $(ii)$ with synonymous substitutions, to remove positively selected variants. Here, we considered a 300-base window, which was iteratively slid along the genome 1 base at a time. For every iteration and for each window, we first computed the total number of distinct synonymous SNVs fall within the window, and normalized with respect to the window length. Finally, we associate a density value to every genome location, that is the mean value of all windows which include such location.

We executed the analysis on Dataset #1 and we further investigated the properties of: $(i)$ the 5 clonal variants identified as possible homoplasies via VERSO STEP #1, $(ii)$ 80 selected minor variants: (1) detected in multiple clonal genotypes (identified via VERSO STEP #1), (2) detected in at least 10 samples, (3) nonsynonymous, (4) falling on a region of the genome with mutational density lower than the median value ($= 0.083$ [syn. mutations][nucleotides]$^{-1}$) (4 mutations were filtered-out after manual curation).

In Figure S4 the mutational density of the SARS-CoV-2 genome (Fig. S4**A**) and the related empirical distribution (Fig. S4**B**) are shown, as computed from the 2906 samples of Dataset #1, by highlighting the clonal variants mentioned above. Furthermore, Tables S5 and S6 include summary features for all such variants.

Importantly, this analysis allows to assess whether a homoplasy detected via VERSO might have been positively selected due to some functional advantage or, instead, if it may be due to mutational hotspots or phantom mutations. The results on Dataset #1 are discussed in the main text.

**Homoplasies: clonal variants**

| Variant | Genome location | Variant effect | Amino Acid substitution | # of samples | Mutational density |
|---|---|---|---|---|---|
| 11083 G>T | ORF1ab | NS | 3606 L>F | 460 | 0.085 |
| 27964 C>T | ORF8 | NS | 24 S>L | 182 | 0.124 |
| 28311 C>T | N | NS | 13 P>L | 153 | 0.100 |
| 14805 C>T | ORF1ab | S | - | 237 | 0.082 |
| 18555 C>T | ORF1ab | S | - | 152 | 0.103 |

Table S5: **Summary features of candidate homoplastic clonal variants.** The table includes the main features of the 5 clonal variants identified as candidate homoplasies via VERSO STEP #1.

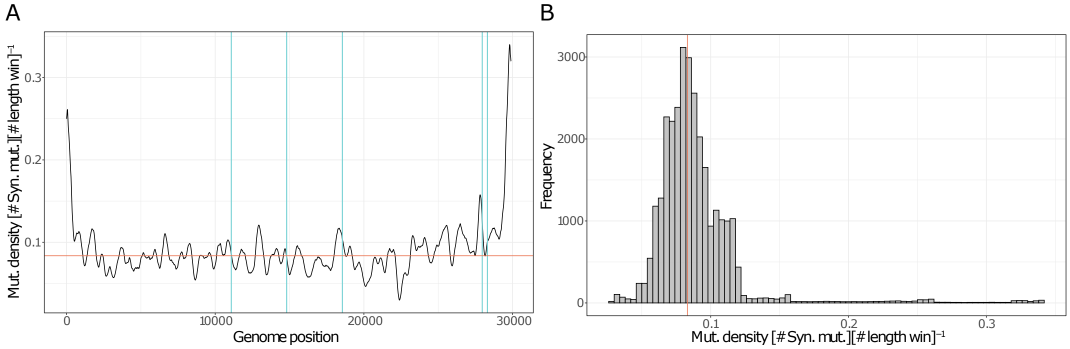


Figure S4: **Characterization of the mutational density over the SARS-CoV-2 genome. (A)** The mutational density, evaluated on synonymous minor variants detected in the samples of Dataset #1 via the sliding-window approach described in the text, is projected on the SARS-CoV-2 genome. The vertical lines represent the positions of 5 clonal variants involved in homoplasies and discussed in the main text (light blue) (the horizontal solid line represent the median mutational density). **(B)** The related empirical distribution is shown.

##### 2.4 Bottleneck size estimation – Dataset #1

To have an estimation of the size of the genetic bottleneck in SARS-CoV-2 we evaluate how the variance of some selected SNVs detected in the samples of Dataset #1 changes upon time[12,13]. In principle, if the evolution is purely neutral, the changes in the genetic variance between two different time points are exclusively due to transmission bottleneck effects.

We first divided the time interval under study (i.e., 30 weeks, from the 1-st week of the 2020 to 29-th week of 2020) into two equal intervals (i.e., $t_1 \leq$ 14-th week or $t_2 >$ 14-th week).

Then, in order to minimize the effect of the genetic drift and to consider comparison between groups in which a transmission event might have occurred, we restrict the analysis to group of samples characterized by distinct clonal genotypes identified via VERSO STEP #1. It is also sound to select a number of SNVs that are presumably neutral or quasi-neutral markers, as proposed in[12,13]. To this end, we employed the following criteria: *i)* we take into account only

| Variant | # of clonal genotypes | # of samples | Type | Genomic location | Amino Acid substitution | Mutational density | Average VF |
|---|---|---|---|---|---|---|---|
| 5743 G>C | 15 | 96 | alway minor | ORF1ab | 1826E>D | 0.08 | 0.159 |
| 5744 T>C | 15 | 95 | alway minor | ORF1ab | 1827Y>H | 0.08 | 0.178 |
| 17747 C>T | 4 | 78 | mixed | ORF1ab | 5828P>L | 0.066 | 0.601 |
| 2192 A>G | 2 | 51 | alway minor | ORF1ab | 643R>G | 0.071 | 0.094 |
| 12164 G>T | 12 | 50 | alway minor | ORF1ab | 3967A>S | 0.069 | 0.181 |
| 2578 T>A | 6 | 48 | alway minor | ORF1ab | 771S>R | 0.065 | 0.095 |
| 24557 G>T | 12 | 47 | alway minor | S | 999G>C | 0.075 | 0.141 |
| 8790 G>T | 11 | 47 | alway minor | ORF1ab | 2842G>V | 0.064 | 0.167 |
| 11535 G>T | 12 | 46 | alway minor | ORF1ab | 3757G>V | 0.076 | 0.15 |
| 17096 G>T | 12 | 46 | alway minor | ORF1ab | 5611G>V | 0.079 | 0.112 |
| 7038 G>T | 11 | 45 | alway minor | ORF1ab | 2258G>V | 0.078 | 0.184 |
| 2036 G>T | 13 | 43 | alway minor | ORF1ab | 591A>S | 0.078 | 0.143 |
| 9249 G>T | 11 | 43 | alway minor | ORF1ab | 2995G>V | 0.08 | 0.135 |
| 13571 G>T | 12 | 40 | alway minor | ORF1ab | 4436G>V | 0.065 | 0.122 |
| 5766 G>C | 12 | 40 | alway minor | ORF1ab | 1834G>A | 0.081 | 0.11 |
| 20135 T>A | 4 | 40 | mixed | ORF1ab | 6624V>E | 0.046 | 0.47 |
| 14906 A>C | 3 | 40 | alway minor | ORF1ab | 4881N>T | 0.069 | 0.068 |
| 5765 G>A | 12 | 39 | alway minor | ORF1ab | 1834G>S | 0.081 | 0.11 |
| 4505 G>T | 11 | 39 | alway minor | ORF1ab | 1414G>C | 0.065 | 0.122 |
| 2604 G>T | 11 | 38 | alway minor | ORF1ab | 780G>V | 0.062 | 0.115 |
| 7214 G>T | 11 | 34 | alway minor | ORF1ab | 2317D>Y | 0.073 | 0.085 |
| 22899 G>T | 10 | 34 | alway minor | S | 446G>V | 0.06 | 0.152 |
| 9141 G>T | 10 | 33 | alway minor | ORF1ab | 2959G>V | 0.083 | 0.152 |
| 17848 G>T | 9 | 33 | alway minor | ORF1ab | 5862G>C | 0.069 | 0.088 |
| 16787 G>T | 11 | 32 | alway minor | ORF1ab | 5508G>V | 0.078 | 0.07 |
| 7246 G>T | 10 | 32 | alway minor | ORF1ab | 2327W>C | 0.073 | 0.103 |
| 15890 C>A | 4 | 28 | alway minor | ORF1ab | 5209T>K | 0.081 | 0.324 |
| 5375 G>T | 9 | 26 | alway minor | ORF1ab | 1704A>S | 0.081 | 0.116 |
| 7017 G>T | 9 | 25 | alway minor | ORF1ab | 2251G>V | 0.078 | 0.106 |
| 20465 A>G | 4 | 25 | mixed | ORF1ab | 6734D>G | 0.058 | 0.516 |
| 20387 A>G | 3 | 25 | mixed | ORF1ab | 6708K>R | 0.055 | 0.248 |
| 17849 G>T | 10 | 24 | alway minor | ORF1ab | 5862G>V | 0.069 | 0.077 |
| 14536 C>T | 7 | 24 | alway minor | ORF1ab | 4758L>F | 0.078 | 0.069 |
| 16514 A>G | 2 | 24 | mixed | ORF1ab | 5417Y>C | 0.064 | 0.24 |
| 3004 G>T | 10 | 23 | alway minor | ORF1ab | 913E>D | 0.064 | 0.118 |
| 8696 G>T | 10 | 22 | alway minor | ORF1ab | 2811A>S | 0.055 | 0.072 |
| 2247 G>T | 9 | 22 | alway minor | ORF1ab | 661G>V | 0.076 | 0.08 |
| 3096 C>T | 12 | 21 | mixed | ORF1ab | 944S>L | 0.058 | 0.204 |
| 16188 G>T | 10 | 21 | alway minor | ORF1ab | 5308W>C | 0.059 | 0.067 |
| 3259 G>T | 9 | 21 | alway minor | ORF1ab | 998Q>H | 0.061 | 0.091 |
| 17380 T>C | 3 | 21 | mixed | ORF1ab | 5706Y>H | 0.07 | 0.28 |
| 16516 A>G | 2 | 21 | mixed | ORF1ab | 5418K>E | 0.065 | 0.252 |
| 7363 G>T | 10 | 20 | alway minor | ORF1ab | 2366W>C | 0.067 | 0.073 |
| 11556 A>T | 3 | 20 | alway minor | ORF1ab | 3764E>V | 0.078 | 0.083 |
| 17929 A>C | 2 | 20 | alway minor | ORF1ab | 5889I>L | 0.077 | 0.286 |
| 9223 C>A | 2 | 20 | alway minor | ORF1ab | 2986H>Q | 0.082 | 0.224 |
| 9843 T>A | 4 | 19 | alway minor | ORF1ab | 3193L>* | 0.07 | 0.122 |
| 17933 C>G | 2 | 19 | alway minor | ORF1ab | 5890T>S | 0.077 | 0.241 |
| 12141 C>T | 10 | 18 | alway minor | ORF1ab | 3959T>I | 0.071 | 0.083 |
| 5617 G>T | 9 | 18 | alway minor | ORF1ab | 1784Q>H | 0.079 | 0.068 |
| 15202 G>T | 9 | 18 | alway minor | ORF1ab | 4980V>L | 0.071 | 0.067 |
| 21137 A>G | 8 | 18 | mixed | ORF1ab | 6958K>R | 0.081 | 0.227 |
| 15157 C>A | 8 | 18 | alway minor | ORF1ab | 4965Q>K | 0.069 | 0.191 |
| 17663 A>G | 2 | 18 | alway minor | ORF1ab | 5800Y>C | 0.069 | 0.157 |
| 19252 G>T | 7 | 17 | alway minor | ORF1ab | 6330V>L | 0.079 | 0.078 |
| 16376 C>T | 4 | 16 | mixed | ORF1ab | 5371P>L | 0.058 | 0.327 |
| 11562 G>A | 2 | 16 | alway minor | ORF1ab | 3766C>Y | 0.079 | 0.193 |
| 4917 T>A | 2 | 16 | mixed | ORF1ab | 1551I>N | 0.071 | 0.147 |
| 16508 G>T | 8 | 15 | alway minor | ORF1ab | 5415G>V | 0.064 | 0.081 |
| 2891 G>A | 2 | 15 | mixed | ORF1ab | 876A>T | 0.07 | 0.635 |
| 5658 T>A | 2 | 15 | mixed | ORF1ab | 1798V>E | 0.08 | 0.221 |
| 15199 G>T | 8 | 14 | alway minor | ORF1ab | 4979V>L | 0.071 | 0.067 |
| 5378 G>T | 8 | 14 | alway minor | ORF1ab | 1705G>C | 0.081 | 0.061 |
| 3896 G>T | 7 | 14 | alway minor | ORF1ab | 1211V>F | 0.082 | 0.063 |
| 12053 C>G | 2 | 14 | alway minor | ORF1ab | 3930L>V | 0.08 | 0.197 |
| 12041 G>T | 8 | 13 | alway minor | ORF1ab | 3926D>Y | 0.081 | 0.068 |
| 2250 G>T | 8 | 13 | alway minor | ORF1ab | 662G>V | 0.077 | 0.067 |
| 4854 G>T | 7 | 13 | alway minor | ORF1ab | 1530R>I | 0.068 | 0.092 |
| 4746 C>T | 2 | 13 | alway minor | ORF1ab | 1494S>F | 0.068 | 0.144 |
| 17365 T>C | 2 | 13 | alway minor | ORF1ab | 5701S>P | 0.071 | 0.112 |
| 16068 G>T | 8 | 12 | alway minor | ORF1ab | 5268E>D | 0.064 | 0.081 |
| 19679 G>T | 6 | 12 | alway minor | ORF1ab | 6472G>V | 0.076 | 0.075 |
| 21301 C>T | 2 | 12 | alway minor | ORF1ab | 7013P>S | 0.07 | 0.115 |
| 25913 G>C | 5 | 11 | alway minor | ORF3a | 174G>A | 0.072 | 0.111 |
| 8688 G>T | 5 | 11 | alway minor | ORF1ab | 2808G>V | 0.055 | 0.082 |
| 11288 T>G | 2 | 11 | alway minor | ORF1ab | 3675S>A | 0.069 | 0.13 |
| 16417 A>G | 2 | 11 | alway minor | ORF1ab | 5385T>A | 0.059 | 0.098 |
| 13627 G>T | 7 | 10 | mixed | ORF1ab | 4455D>Y | 0.073 | 0.219 |
| 12079 G>T | 3 | 10 | mixed | ORF1ab | 3938R>S | 0.078 | 0.183 |
| 24552 T>C | 2 | 10 | alway minor | S | 997I>T | 0.075 | 0.092 |

Table S6: **Summary features of candidate homoplastic minor variants.** The table includes the main features of the 80 minor variants identified as candidate homoplasies, ordered by number of samples in which they are detected (see the main text).

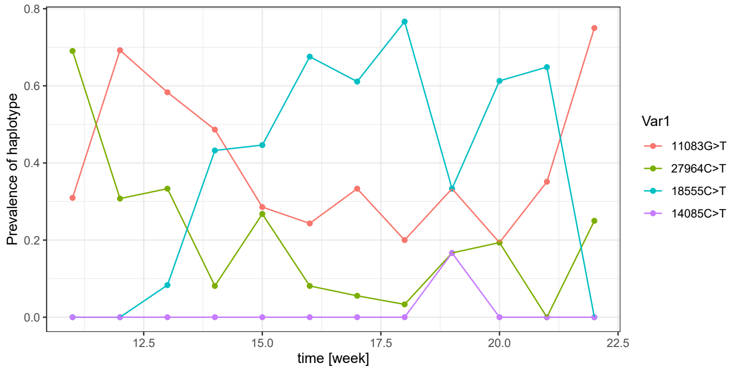


Figure S5: **Dynamics of the haplotypes related to $4$ candidate homoplastic clonal variants.** Temporal evolution of prevalence of $4$ haplotypes associated to homoplastic clonal variants discussed in the main text. The prevalence of the SNV g.28311C>T is not shown, because related samples have no collection date.

*always minor* variants (i.e., variants with VF $< 90\%$ in all samples in which they are detected), *ii)* the SNV must either be a synonymous change at the third codon position or fall in a non-coding region, *(iii)* the SNV must be observed in at least 3 samples in each time interval and, *iv)* no significant changes in mean VF of the SNV must be observed between the two time intervals (i.e., $\leq 0.10$). Notice that the analysis is performed on groups of samples with distinct clonal genotypes, separately.

Applying these filters, we identified 3 distinct SNVs as quasi-neutral markers for Dataset $\#1$ (see Table S7 for the complete list of markers), two them (g.634T>C and g.14523A>G) are detected in genotype G15, while g.15168G>A is detected in genotypes G17 and G21. Note that only the samples with collection date were used in the analysis.

In Figure S6, the distribution of the VF of all markers, divided by time point and clonal genotype is presented. From this analysis, the presence of mild bottleneck effects is highlighted, as confirmed in other study considering the donor-host data[14].

| Mutation id. | VF Mean at $t_1$ | VF Mean at $t_2$ | N. Obs. $t_1$ | N. Obs. $t_2$ | Genotype |
|---|---|---|---|---|---|
| g.634T>C | $0.141 \pm 0.005$ | $0.082 \pm 0.001$ | 10 | 10 | G15 |
| g.14523A>G | $0.253 \pm 0.017$ | $0.235 \pm 0.015$ | 14 | 17 | G15 |
| g.15168G>A | $0.10 \pm 0.0005$ | $0.071 \pm 0.001$ | 4 | 4 | G17 |
| g.15168G>A | $0.152 \pm 0.003$ | $0.091 \pm 0.00005$ | 25 | 5 | G21 |

Table S7: **Summary features of the three SNVs selected as neutral markers employed in the bottleneck size estimation analysis.**

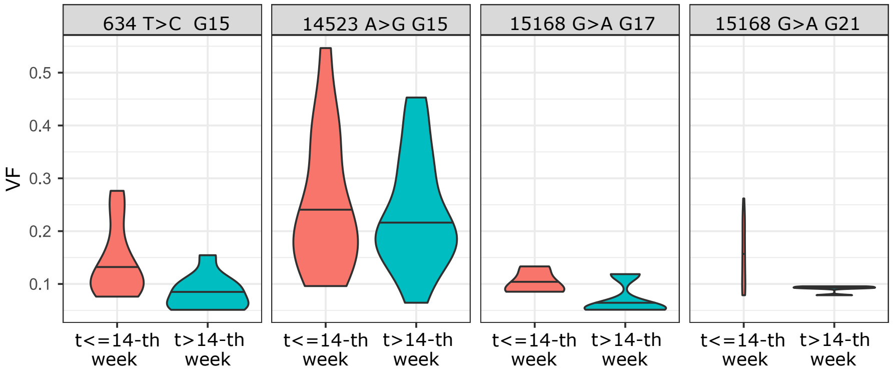


Figure S6: **Bottleneck size estimation** . For each SNV selected as neutral or quasi neutral marker (see Table S7) we show the violin plot of the VF distribution with respect to each clonal genotype, in the subsequent time intervals $t \leq 14$ and $t > 14$ .

##### 2.5 Impact of clonal variant threshold on VERSO stability – Dataset $\#1$

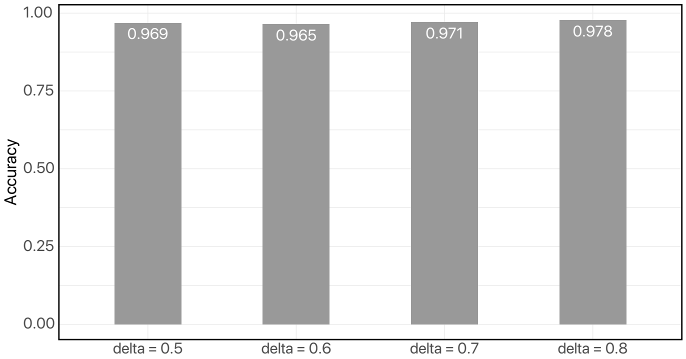


Figure S7: **Stability of VERSO to clonal variant threshold.** Variation of the tree accuracy obtained by changing the threshold used to define a clonal variants ($\delta$), by setting as ground-truth the model returned by VERSO STEP $\#1$ with $\delta = 0.9$.

We assessed the robustness of the results produced by VERSO STEP $\#1$ on Dataset $\#1$ when different thresholds are employed to call clonal variants. In the analysis presented in the main text the default VF threshold ($\delta = 90\%$) was employed, which resulted in $m = 29$ clonal variants.

Here, we scanned a number of thresholds in the set $\delta \in \{50\%, 60\%, 70\%, 80\%\}$, which resulted in 31, 31, 30 and 30 variants, respectively. We then assessed the robustness of VERSO STEP $\#1$ computing the accuracy: $\frac{\text{TP+TN}}{\text{TP+TN+FP+FN}}$ by considering as ground-truth the topology returned by VERSO with $\delta = 90\%$. In this case, the true positives (TP) are the rightly inferred edges, the false positives (FP) the wrongly inferred edges, the false negatives (FN) the edges that are present in the ground-truth topology, but are not inferred, the true negatives (TN) the edges that are not inferred and are not present in the ground-truth topology.

As one can see in Figure S7, the accuracy varies between $0.97$ and $0.98$ in all settings, demonstrating the the results produced by VERSO STEP $\#1$ are robust with regard to the choice of the VF threshold for clonal variant identification.

##### 2.6 Application of VERSO to $1438$ samples from RNA sequencing data (Dataset $\#2$)

The application of VERSO to Dataset $\#2$ is discussed in the main text and the output models are displayed in Figure S8 (see Table S3 for further details).

##### 2.7 Expanded clonal variants tree – Dataset $\#2$

In Figure S9 we report the *expanded* clonal variants tree of Dataset $\#2$ (see above).

##### 2.8 Computation time

We assessed the computation time of VERSO STEP $\#1$ in different simulated scenarios, by especially evaluating the impact of variations of the number of samples $n$. All simulations were performed by employing a specific simulation setting with $m = 14$ clonal variants (simulation id: $\#1$) and we assessed the computation time by varying the number of samples: $n \in \{100, 500, 1000, 5000, 10000\}$. For datasets with more than $1000$ samples, random resampling of the original dataset was performed.

For each configuration, we executed $10$ independent computations, with $1000$ MCMC iterations and $5$ restarts, on a single core of a MacBook Pro, processor 2.6 GHz Intel Core i7, 16 GB RAM 2400 MHz DDR4. The distribution of the run times with respect to the distinct configurations is shown in Supplementary Fig. S10.

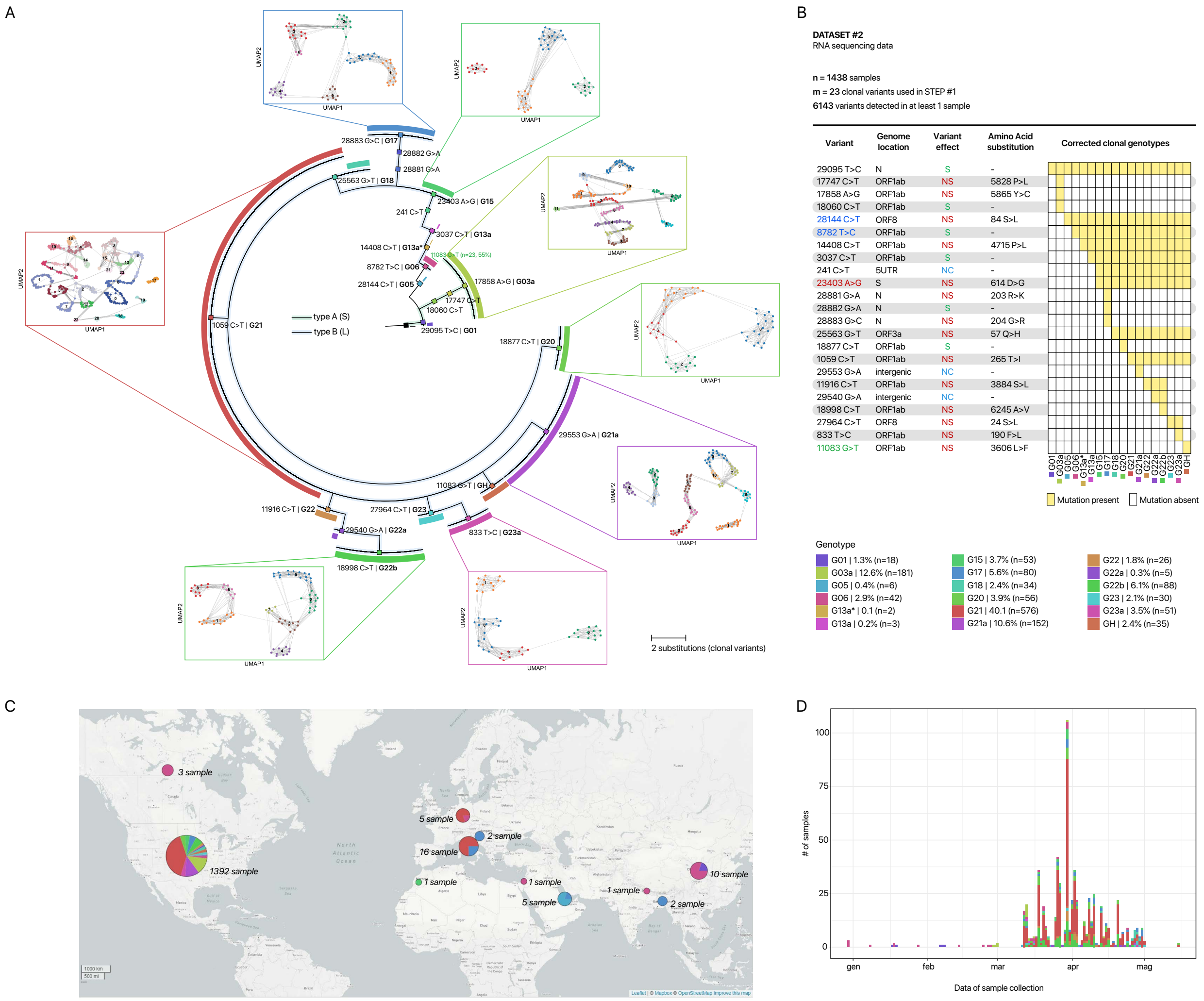


Figure S8: **Viral evolution and intra-host genomic characterization of** 1438 **SARS-CoV-2 samples via VERSO (Dataset** #2**). (A)** The phylogetic model returned by VERSO STEP #1 from the mutational profile of 1438 samples selected after QC, on 23 clonal variants (VF $>$ 90%) detected in at least 3% of the samples of Dataset #2 (reference genome: SARS-CoV-2-ANC). 18 distinct clonal genotypes are identified by VERSO, 11 of which are identical to those found in the analysis of Dataset #1 and maintain the same label. 6 further clonal genotypes are distinct, but evolutionary consistent and are marked with letters $a$, $a*$ and $b$, while clonal genotype GH is associated to homoplastic clonal variant g.11083G>T (the mapping with the lineage nomenclature proposed in[15] is provided in Supplementary File 3). Samples with identical corrected clonal genotypes are grouped in polytomies (visualization via FigTree[16]). Notice that the colors do not correspond with those of Figure 3 of the main text. The green curves juxtaposed to certain polytomies report the number and fraction of samples in which homoplastic mutation g.11083G>T is observed (only if the mutation is detected in at least 10 samples with the same corrected clonal genotype; see Supplementary File 2 for a summary on the samples exhibiting homoplastic clonal variants). The projection of the intra-host genomic composition and similarity computed by VERSO STEP #2 from VF profiles is shown on the UMAP low-dimensional space for the clonal genotypes including $\geq$ 100 samples. Samples are clustered via Leiden algorithm on the k-nearest neighbor graph ($k = 10$), computed on the Bray–Curtis dissimilarity on VF profiles, after PCA. Solid lines represent the edges of the k-NNG. **(B)** The composition of the corrected clonal genotypes returned by VERSO STEP #1 is shown. Clonal SNVs are annotated with mapping on ORFs, synonymous (S), nonsynonymous (NS) and non-coding (NC) states, and related amino acid substitutions (asterisks label nonsense mutations). Variants colored in blue characterize the B (L) type[17], variant g.23403 A>G ($S$, p.614 D>G) is colored in red, homoplastic variant g.11083G>T ($ORF1ab$, p.3606L>F) in green. **(C-D)** The geo-temporal localization of the clonal genotypes via Microreact[18] and the prevalence variation in time are displayed.

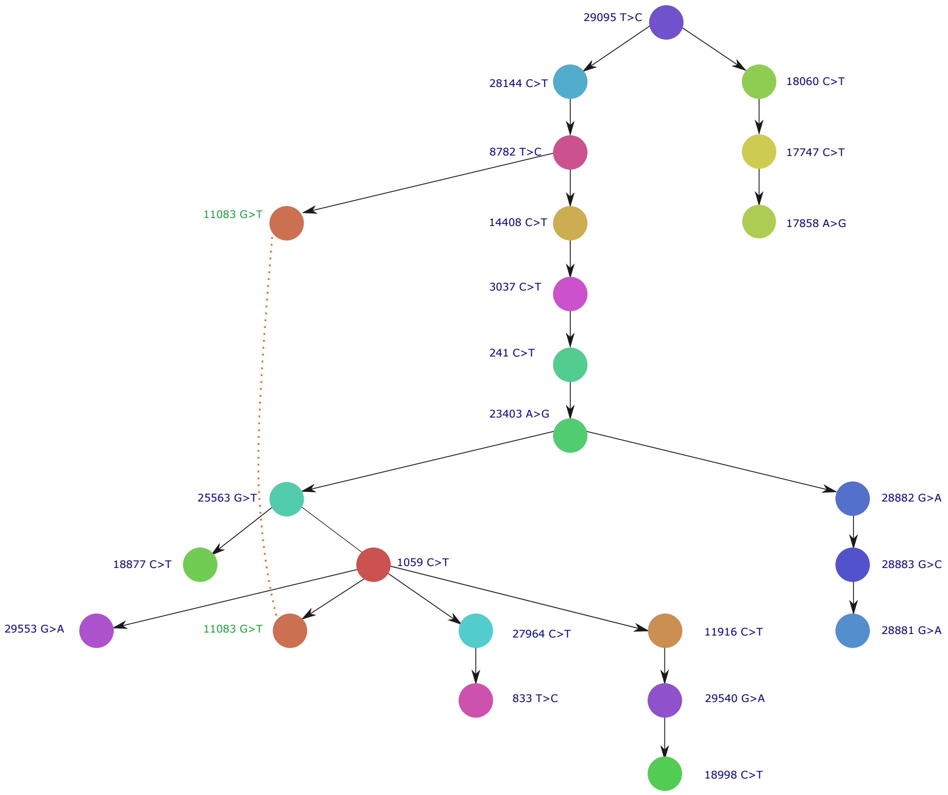


Figure S9: **Expanded clonal variants tree – Dataset #2.** The clonal variants tree **B** representing the inner structure of the output phylogenetic tree inferred from Dataset #2 via VERSO STEP #1 is expanded by duplicating homoplastic variants. Nodes represent clonal variants (consistent colors with Supplementary Figure 8), whereas edges indicate temporal ordering/accumulation relations. Homoplastic mutations were duplicated if observed in at least 10 samples with the same corrected clonal genotype returned by VERSO STEP #1 and are attached after the corresponding characterizing mutation. Homoplastic variants observed in multiple samples with the same corrected clonal genotype are ordered by the number of samples in which they are detected. Dashed lines connect identical homoplastic mutations.

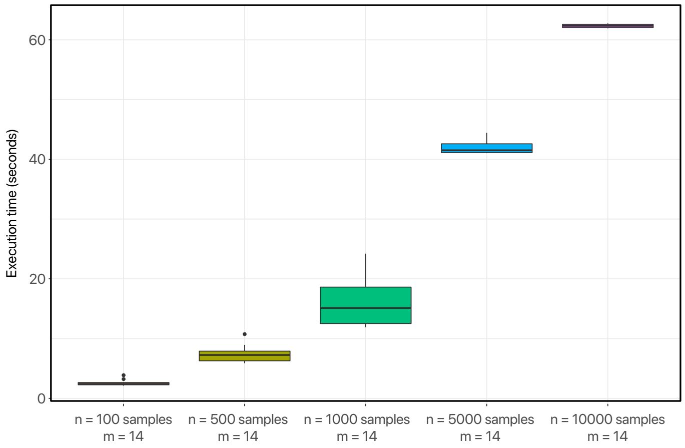


Figure S10: **Computation time**. Computation time of VERSO with respect to the 6 different configurations described in the text; every boxplot displays the distribution of 10 independent computations with 1000 MCMC iterations and 5 restarts. Execution time in seconds.

#### References


[1] Daniele Ramazzotti, Fabrizio Angaroni, Davide Maspero, Gianluca Ascolani, Isabella Castiglioni, Rocco Piazza, Marco Antoniotti, and Alex Graudenzi. Longitudinal cancer evolution from single cells. *bioRxiv*, 2020.

[14] Alexandra Popa, Jakob-Wendelin Genger, Michael D. Nicholson, Thomas Penz, Daniela Schmid, Stephan W. Aberle, Benedikt Agerer, Alexander Lercher, Lukas Endler, Henrique Colaço, Mark Smyth, Michael Schuster, Miguel L. Grau, Francisco Martínez-Jiménez, Oriol Pich, Wegene Borena, Erich Pawelka, Zsofia Keszei, Martin Senekowitsch, Jan Laine, Judith H. Aberle, Monika Redlberger-Fritz, Mario Karolyi, Alexander Zoufaly, Sabine Maritschnik, Martin Borkovec, Peter Hufnagl, Manfred Nairz, Günter Weiss, Michael T. Wolfinger, Dorothee von Laer, Giulio Superti-Furga, Nuria Lopez-Bigas, Elisabeth Puchhammer-Stöckl, Franz Allerberger, Franziska Michor, Christoph Bock, and Andreas Bergthaler. Genomic epidemiology of superspreading events in austria reveals mutational dynamics and transmission properties of SARS-CoV-2. *Science Translational Medicine*, 2020.

[15] Andrew Rambaut, Edward C. Holmes, Áine O'Toole, Verity Hill, John T. McCrone, Christopher Ruis, Louis du Plessis, and Oliver G. Pybus. A dynamic nomenclature proposal for sars-cov-2 lineages to assist genomic epidemiology. *Nature Microbiology*, 5(11):1403–1407, 2020.

[16] A Rambaut. Figtree v1. 3.1. *http://tree. bio. ed. ac. uk/software/figtree/*, 2009.

[17] Peter Forster, Lucy Forster, Colin Renfrew, and Michael Forster. Phylogenetic network analysis of SARS-CoV-2 genomes. *Proceedings of the National Academy of Sciences*, 2020.

[18] Silvia Argimón, Khalil Abudahab, Richard JE Goater, Artemij Fedosejev, Jyothish Bhai, Corinna Glasner, Edward J Feil, Matthew TG Holden, Corin A Yeats, Hajo Grundmann, et al. Microreact: visualizing and sharing data for genomic epidemiology and phylogeography. *Microbial genomics*, 2(11), 2016.
